# Widespread Utilization of Diverse Organophosphate Pollutants by Marine Bacteria

**DOI:** 10.1101/2021.10.18.464763

**Authors:** Dragana Despotović, Einav Aharon, Olena Trofimyuk, Artem Dubovetskyi, Kesava Phaneendra Cherukuri, Yacov Ashani, Haim Leader, Andrea Castelli, Laura Fumagalli, Alon Savidor, Yishai Levin, Liam M. Longo, Einat Segev, Dan S. Tawfik

## Abstract

Anthropogenic organophosphates (AOPs), such as phosphotriesters, are used extensively as plasticizers, flame retardants, nerve agents and pesticides. Soil bacteria bearing a phosphotriesterase (PTE) can degrade AOPs, but whether bacteria are capable of utilizing AOPs as a phosphorus source, and how widespread PTEs are in nature, remains unclear. Here, we report the utilization of diverse AOPs by four model marine bacteria and seventeen bacterial isolates from seawater samples. To unravel the details of AOP utilization, two novel PTEs from marine bacteria were isolated and characterized. When expressed in *E. coli*, these PTEs enabled growth on a pesticide analog as the sole phosphorus source. Utilization of AOPs provides bacteria with a source of phosphorus in depleted environments and offers a new prospect for the bioremediation of a pervasive class of anthropogenic pollutants.

**One sentence summary:** Widespread utilization of diverse organophosphate pollutants by over 20 marine bacterial strains represents a new hope for ocean bioremediation.

## Main text

Phosphorus (P) is an essential component of many key biological metabolites and is thus competitively utilized, particularly in marine environments, which can be P limited *(1,2)*. Microbial uptake of P preferentially occurs through phosphate or phosphate-containing natural metabolites – mainly phosphomonoesters, but also phosphodiesters and, to a lesser extent, phosphonates and phosphites. However, not all microorganisms can utilize all phosphorus sources *(2–4)* because multiple, specialized enzymes are required (**Fig. 1A**). With the advent of industrialization, modern environments now have an additional source of P-containing compounds: anthropogenic organophosphates (AOPs), which includes phosphotriesters as well as diesters of both phosphonates and phosphites *(5)*. These man-made compounds are used in an array of applications; phosphotriesters are pesticides *(6)*, flame retardants *(7)* and plasticizers *(8)*, while phosphonate diesters can act as potent nerve agents *(9)*. Over the last several decades, the concentration of AOPs in the ocean has been steadily increasing *(10,11)* and can potentially exceed the concentration of phosphate in P-depleted environments *(1,10)* raising the question: Are AOPs a viable alternative P source in environments with limiting concentrations of labile phosphate?

**Fig. 1:**
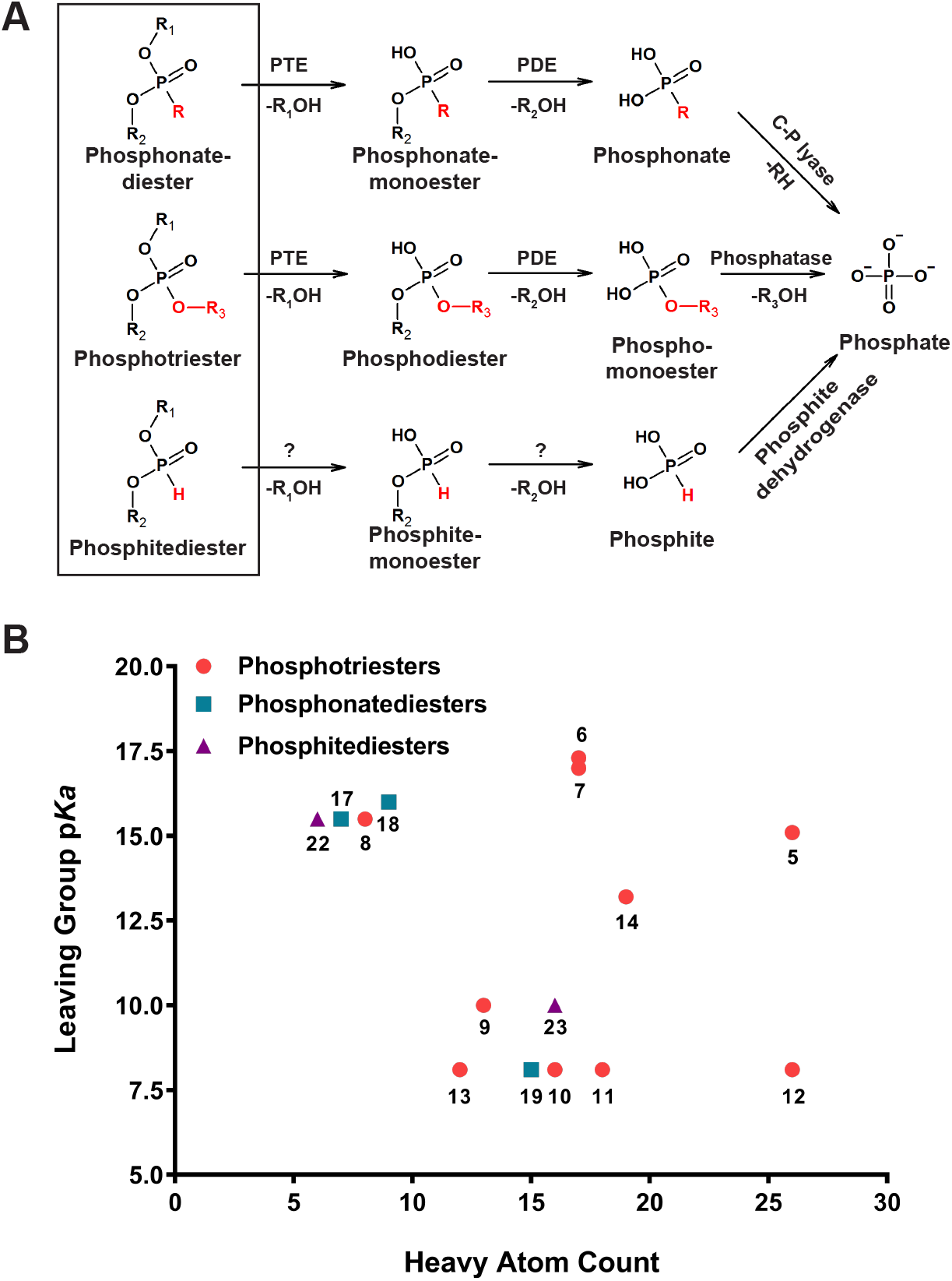
Anthropogenic organophosphate (AOP) bioremediation pathway and the distribution of AOP chemical properties used in this study. **A** Anthropogenic organophosphates, black rectangle, include phosphotriesters and their phosphonate and phosphite diester analogs. AOPs share a similar structure with the exception of the groups colored in red. Phosphotriesterases (PTEs) hydrolyze AOPs to phosphodiesters and phosphonate monoesters, which are further hydrolyzed by phosphodiesterases (PDEs). Enzymatic degradation of phosphitediesters is not yet well described, though is suspected to use PTEs and PDEs. The last step in the AOP utilization pathway, conversion to phosphate, is mediated by three different enzymes: alkaline phosphatases hydrolyze phosphomonoesters to phosphate, C-P lyase converts phosphonates to phosphate, and phosphite dehydrogenase converts phosphite to phosphate. **B** The distribution of substrate properties used for growth profiling of marine bacteria. Each compound number can be found in **Table S1** with its structural representation. These compounds are analogs of flame retardants, plasticizers, nerve agents and pesticides. The compounds vary greatly in both the leaving group potential (shown here is the p*Ka* of the best leaving group) and the overall size of the compound.

As a potential nutrient for bacteria, AOPs have some favorable properties: First, they are very hydrophobic and can readily diffuse across membranes so, unlike charged phosphates, they are unlikely to need dedicated transporters. Second, natural P sources are competitively utilized; thus, if AOP utilization is a specialized trait, it may confer a fitness advantage. Unlike phosphodiesterases (PDEs) and phosphatases, phosphotriesterases (PTEs) – which convert AOPs into metabolically labile mono- and di-esters – have been detected in only a handful of soil bacteria that degrade paraoxon, a common AOP pollutant *(12,13)*. Several recent reports have noted indications of microbial utilization of AOPs in seawater – such as stimulated growth upon addition of AOP flame retardants *(14,15)*; PTE activity when the growth medium was supplemented with methyl paraoxon *(15)*; and the observation of an alkaline phosphatase from *Alteromonas mediterranea* that can promiscuously act as a PTE *(16)* – but it still remains unclear whether AOPs, in the absence of another P source, can support microbial growth. To address this question, and uncover the biochemical details of AOP metabolism, we have systematically assayed the growth potential of model marine bacteria and environmental isolates on media containing a variety of AOPs as the sole P source.

The chemical properties of the R-groups in AOPs are diverse and tuned for specific applications. For example, pesticides and nerve agents are often designed with good leaving groups – that is, conjugate acids with a low p*Ka* – while plasticizers and flame retardants, on the other hand, often bear poor leaving groups. As for R-group size, flame retardants, in particular, range from bulky to compact, depending on the industrial process. Phosphotriester derivatives are also commonly used: thiophosphates (P=S) and alkyl thiols (P-S-alkyl) are pesticides (and nerve agents) while phosphite diesters stabilize plastics. Thus, to broadly sample the chemical space of modern AOPs as a potential P source, we synthesized 15 representative AOP and AOP analogs with diverse chemical properties (**Fig. 1B, Table S1**).

To estimate the extent of AOP degradation and metabolism that can be realized by marine bacteria, environmental samples were collected at (i) the nature reserve in the Gulf of Eilat, Red Sea, Israel, and (ii) three locations in the Mediterranean Sea, Israel. The Mediterranean Sea is particularly interesting because phosphate concentrations drop below 10 nM during the summer, which is below the uptake affinity of many organisms *(17)*. The seawater samples were used to inoculate artificial seawater media containing AOPs as the only P source and with various defined carbon sources (**Fig. 2**). Remarkably, all cultures exhibited growth on all tested AOPs (compounds 8, 9, 10, 12 in **Table S1** and 36 in **Table S2**, selected to have a range of chemical properties). Passages of the cultures in liquid and solid media resulted in the isolation of 17 bacterial strains from 5 orders of proteobacteria (**Fig. 2 and Table S3)**. The isolated strains are known to be abundant in the Mediterranean Sea *(18)* and the Red Sea *(19)* – revealing that common bacteria in these environments are able to utilize several AOPs as the sole source of P.

**Fig. 2:**
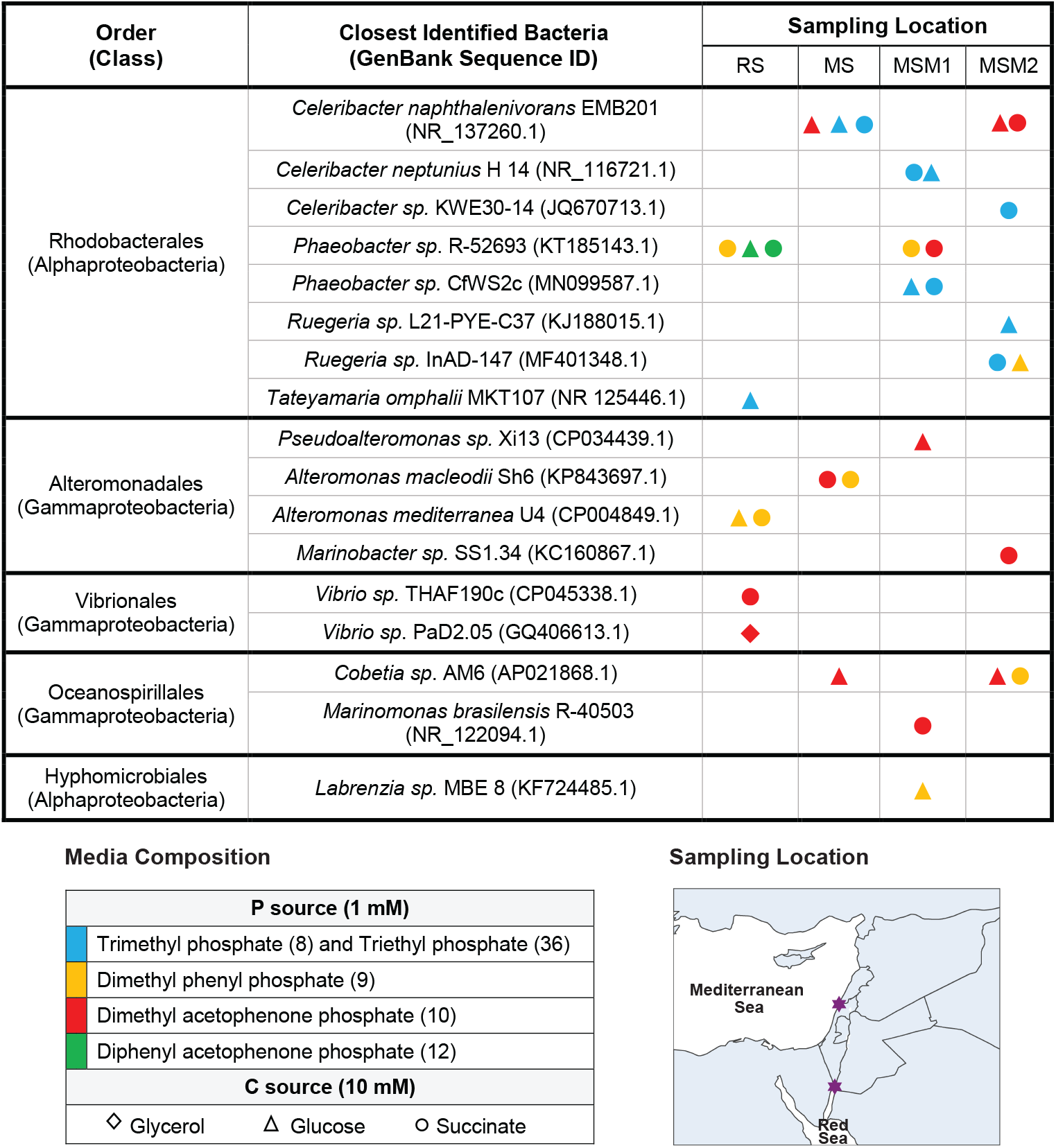
Isolation and identification of marine bacteria that grow on organophosphates. Bacterial strains were isolated from the Red Sea and the Mediterranean Sea based on their ability to grow on various organophosphates as the sole P source (indicated in different colors). Different carbon sources were tested to maximize the number of strains isolated and included glucose (diamonds), succinate (triangles) and glycerol (circles). RS – Red Sea; MS – Mediterranean Sea, 600 m from the shore; MSM1 - Mediterranean Sea, coastal water mixed with fresh water at location 1; MSM2 - Mediterranean Sea, coastal water mixed with fresh water at location 2. SLocations are specified in the Materials and Methods section.

To better characterize the ability of diverse AOP compounds to support the growth of marine bacteria, 10 species from the isolated environmental strains (selected to maximize diversity) and four well-characterized marine bacteria – *Ruegeria pomeroyi* DSS3 *(20), Ruegeria sp*. TM1040 *(21), Phaeobacter inhibens (22)* and *Dinoroseobacter shibae (23)* – were assayed for growth on 15 AOPs and AOP analogs as the sole P source (**Fig. 3** and **Figs. S1-S15**). The resulting growth profiles further confirmed the ability of multiple bacterial strains to use a variety of AOPs while the patterns of AOP utilization revealed two general trends: First, better leaving group potentials support more robust bacterial growth (*e*.*g*., 4-hydroxy acetophenone); and second, almost all sources of P that were tested – with the exception of tributyl phosphate (compound 7) – were metabolically accessible to the tested bacteria. Even non-activated AOPs, such as tris(2-butoxyethyl) phosphate (compound 5, a widely used flame retardant in plastics and rubbers) could be harnessed as a source of P by 5 different species in the absence of alternatives. The failure of *E. coli* to grow on any phosphotriesters serves as a reminder that PTE activity is not trivially present in all bacteria – in fact, it is quite rare *(12,16,24,25)* – potentially suggesting unique enrichment of PTEs in marine ecosystems. Finally, the ability of about half of the tested bacterial species to grow on phosphonate diesters (commonly used as nerve agents) indicates a reasonably wide distribution of C-P lyases, which convert methyl phosphonic acid to inorganic phosphate (**Fig. 1A**). Using the most similar bacterial strains with a sequenced genome as a proxy (see **Fig. 2** and **Table S3** for more details), we searched for homologs of PhnJ, the catalytic core of the C-P lyase pathway in *E. coli*; indeed, strains associated with growth on phosphonate diesters were uniquely identified to contain a homolog of PhnJ.

**Fig. 3:**
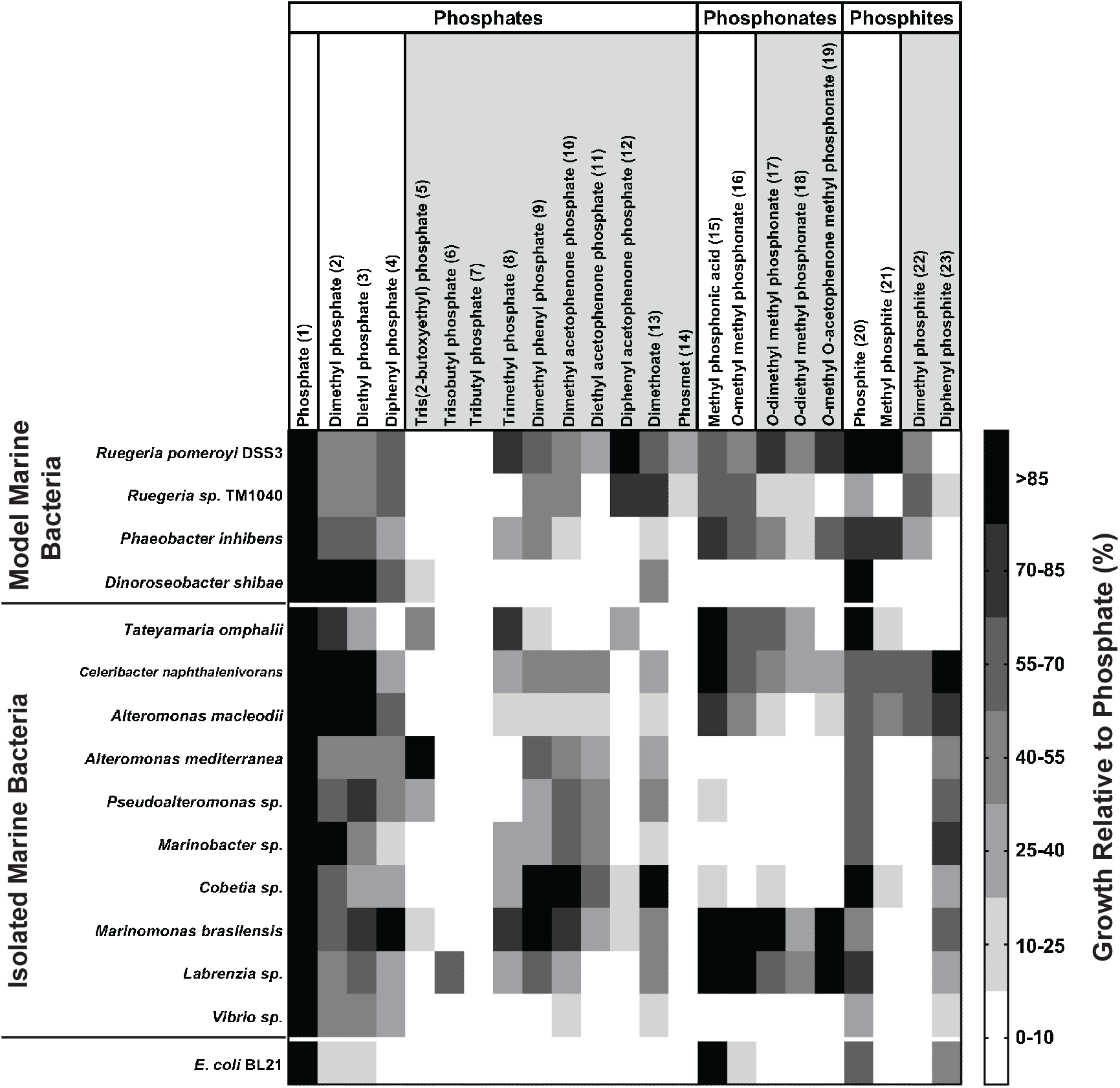
Growth potential of model marine bacteria and environmental isolates on selected organophosphates. Each compound number is in brackets and can be found in **Table S1** with a full structural representation or in **Figure 1B** for a more general description of its chemical properties. Anthropogenic chemicals – that is, phosphotriesters and their analogs phosphonate and phosphite diesters – are shaded in light gray. The growth potentials on each substrate were calculated from the maximum optical density obtained during growth on the substrate OP normalized by the maximal optical density obtained during growth on phosphate. See **Figures S1-S15** for the raw growth curves.

To understand the mechanics of AOP utilization more generally, we endeavored to identify which enzymes enabled growth when AOPs are the sole source of P. While phosphatases and PDEs are widely distributed, PTEs are rare and often only hydrolyze phosphotriesters with activated leaving groups like p-nitrophenol (pNP) *(12,16,24,26)*. We thus attempted to isolate PTEs from the two *Ruegeria* species since they exhibited robust growth on a variety of AOPs but had different substrate preferences (**Fig. S16**). Using PTE activity against chromogenic AOP analogs, *in situ* activity gels and shot-gun proteomics (**Table S2, Fig. S17**; see **Supplemental Information** for more details), two candidate genes encoding PTEs were identified. Both genes encode a periplasmic protein, which – given the proximity to the outer membrane – is consistent with the utilization of compounds present in the extracellular environment. Next, a gene abundance analysis of the TARA dataset found that these enzymes are common in the marine environment and present in a significant fraction of the bacterial community (**Fig. S18)**. Both of the purified gene products were confirmed to be active PTEs against multiple phosphotriesters (**Fig. 4A, Table S4**) with second-order rate constants similar to that of the average enzyme in the BRENDA database of enzyme constants *(27)*.

**Fig. 4:**
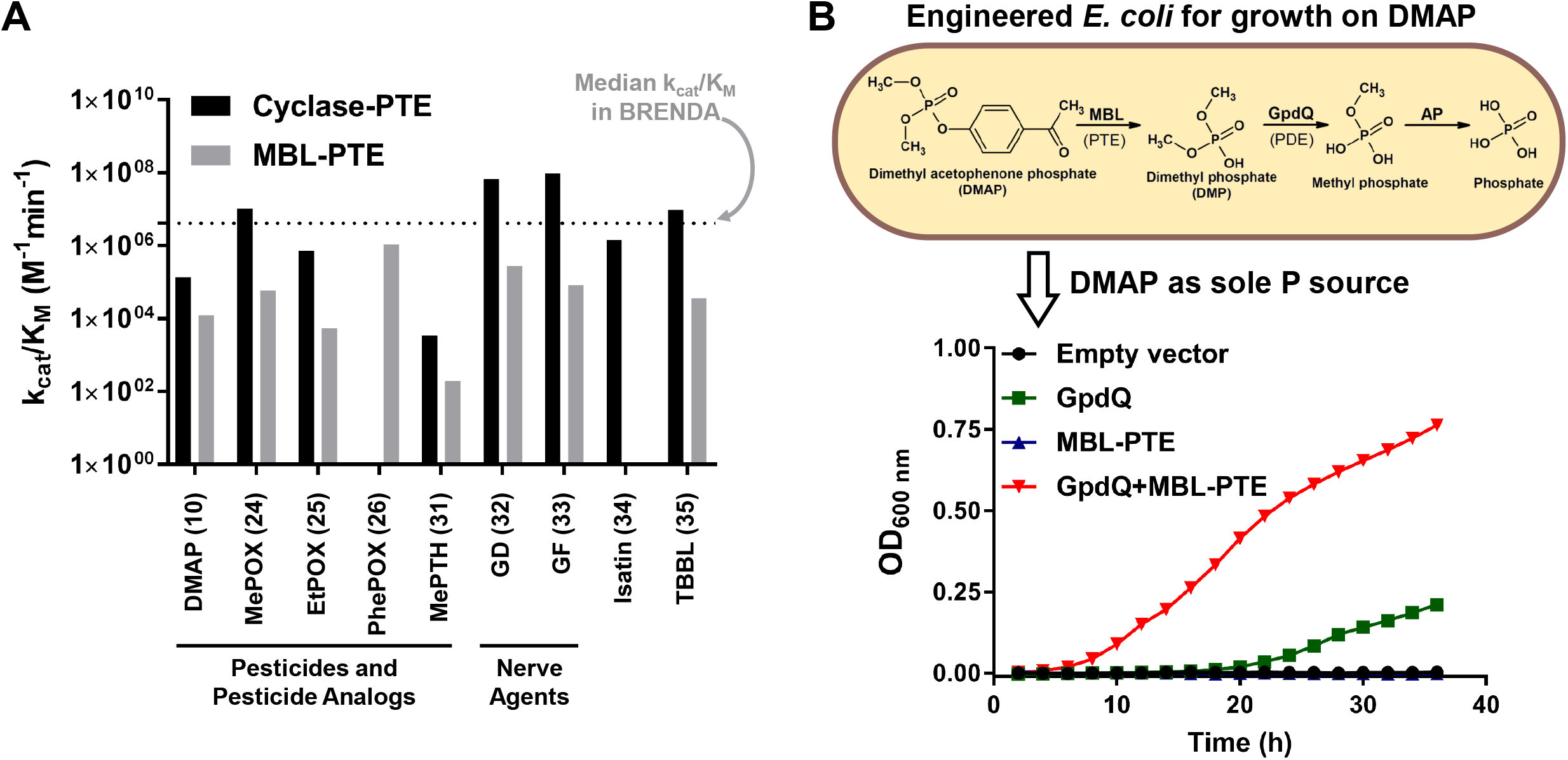
Kinetic characterization of the newly discovered PTEs and engineering *E. coli* to utilize an AOP as the sole P source. **A** The catalytic efficiency (k_cat_/K_M_) of cyclase-PTE from *R. pomeroyi* DSS3 and the metallo-beta lactamase (MBL-PTE) from *Ruegeria sp*. TM1040 were determined for various AOPs (pesticides, their analogs and nerve agents). Catalytic efficiency for the activity of the closest structural homologues was also measured (isatin for IH activity and TBBL for lactonase activity). The dotted line indicates the median catalytic efficiency (k_cat_/K_M_ = 4.2×10^6^ M^-1^min^-1^) of the enzymes available in BRENDA database. **B** Growth of *E. coli* BL21 co-expressing GpdQ and MBL-PTE in media with dimethyl acetophenone phosphate (DMAP, compound 10) as the sole P source. Only co-expression of a phosphotriesterase (MBL-PTE) and a phosphodiesterase (GpdQ) resulted in significant, evident growth on DMAP. Expression of only GpdQ supported slight growth, in accordance with the weak phosphotriesterase activity of this enzyme.

Although both PTEs originated from bacterial strains of the same genus, the enzymes themselves are non-homologous, with one enzyme belonging to the metallo-beta-lactamase superfamily (hereafter, “MBL-PTE”; NCBI Reference Sequence: WP_011538812.1) and the other enzyme belonging to the cyclase family (hereafter, “cyclase-PTE”; NCBI Reference Sequence: WP_011046510.1), identified in *Ruegeria sp*. TM1040 and *Ruegeria pomeroyi* DSS3, respectively. Neither enzyme has significant phosphatase or PDE activity and both enzymes efficiently degrade lactones, as is common for PTEs *(26,28,29)* (**Fig. 4A, Fig. S19, Fig. S20** and **Table S4**). While PTE evolution within the metallo-beta-lactamase superfamily has been previously observed, this is the first identified example of PTE emergence within the cyclase family.

The closest known structural homolog of cyclase-PTE is isatin hydrolase (IH) *(30)* and, indeed, the enzyme retains some IH activity, though at a lower level than the PTE activity against most of the tested phosphotriesters and phosphonate analogs (**Fig. 4A, Fig. S20A-E, Table S4)**. IH enzymes, however, do not generally exhibit PTE activity (**Fig. S21)**, suggesting that the *R. pomeroyi* DSS3 enzyme was re-specialized as a PTE. The re-specialization event seems to have involved a change in the catalytic metal, converting the enzyme from a Mn^2+^-dependent IH to a Zn^2+^-dependent PTE (**Fig. S22B)** *(30)*. Cyclase-PTE is also notable because, to the best of our knowledge, this is the first wild-type enzyme that demonstrates high activity against the nerve agents soman (GD; compound 32 in **Table S2**) and cyclosarin (GF; compound 33 in **Table S2**). Moreover, while previously discovered PTEs prefer the non-toxic *R*_P_ isomer or hydrolyze both the *S*_P_ and the *R*_P_ isomers with similar efficiency *(31,32)*, cyclase-PTE efficiently and preferentially hydrolyzes the neurotoxic *S*_P_ isomer (**Fig. 4A, Fig. S20C, D, Table S4**).

Finally, the kinetic characterization of cyclase-PTE and MBL-PTE suggest that either enzyme, in concert with a PDE (**Fig. 1A**), may be able to confer growth on an AOP or AOP analog (**Fig. 4A, Table S4**). To unambiguously establish the role of the newly identified PTEs in bacterial AOP utilization – and determine if PTE functionality can be readily incorporated into the metabolic repertoire of bacteria for growth – we co-expressed in *E. coli* each of the two PTEs with the known PDE GpdQ from *E. aerogenes (33)*. Cyclase-PTE could not be efficiently expressed in *E. coli*, resulting in cell toxicity (**Table S5** and **Fig. S23)**. However, *E. coli* co-expressing MBL-PTE and the PDE GpdQ readily grew on media with the pesticide analog dimethyl acetophenone phosphate (DMAP, compound 10 in **Table S1**) as the sole source of P (**Fig. 4B)**.

Ever opportunistic and metabolically versatile, marine bacteria have adapted to use anthropogenic organophosphates as a source of P. The extent of this adaptation is remarkable, as nearly all of the AOPs and AOP analogs tested could be utilized by at least one bacterial strain (**Fig. 3**). Furthermore, the bacterial strains, themselves, are widely distributed – both phyletically, where they span 5 orders (**Fig. 2)**, and geographically (**Fig. 2** and **Fig. S18**). Indeed, even within the *Ruegeria* genus, the enzymes employed represent independent emergences of PTE activity (**Fig. 4A**). Man-made organophosphates have changed the landscape of P utilization in marine ecosystems, and the results presented herein suggest that the full breadth of AOP utilization is underestimated. Taken together, these results offer new prospects for the efficient bioremediation of ocean waters and agricultural runoff.

## Acknowledgments

We gratefully acknowledge financial support from the Defense Threat Reduction Agency (DTRA) of the US Department of Defense (HDTRA1-17-0057). We are grateful to Prof. Miguel Frada (The Hebrew University of Jerusalem) and Prof. Yael Kiro (Weizmann Institute of Science) for collecting water samples from the Red Sea and the Mediterranean Sea; Dr. Michael Etzerodt (Aarhus University) for sharing the isatin hydrolase gene and Prof. Jonathan Todd (University of East Anglia) and Prof. Andrew Johnston (University of East Anglia) for providing us with the *Ruegeria* strains. We acknowledge Roni Beiralas, Dr. Martin Sperfeld and Yemima Duchin Rapp for helping us establish protocols for marine bacteria cultivation and Jagoda Jabłońsk for helping with Figure S23. We acknowledge insightful comments on the manuscript from Prof. Maria Vila-Costa, Prof. Ita Gruic-Sovulj, Prof. Danica Galonić-Fujimori, and Dr. Martin Sperfeld.

## Author Contributions

K.P.C, H.L., A.C. and L.F. synthesized organophosphate substrates. A.S. and Y.L. performed mass spectrometry experiments and analysis. E.A., O.T., A.D. and Y.A. performed experiments. D.D., E.S. and D.S.T conceived, supervised, analyzed experiment; D.D., L.M.L., E.S. and D.S.T. wrote the manuscript with the input of all authors.

## Data and materials availability

All data are available in the main text or the supplementary materials.

## Supplementary Material

