## Supplementary material for "Widespread Utilization of Diverse Organophosphate Pollutants by Marine Bacteria": Methods and Supplemental Figures and Tables

^†^Dan S. Tawfik died on May 4^th^, 2021 during the preparation of this manuscript.

^*^Corresponding authors: Einat Segev

Dragana Despotović

**This PDF file includes:**

Materials and Methods

Figs. S1 to S24

Tables S1 to S5

**Materials and methods**

**Synthesis of OPs**

General Information: All chemicals and solvents were purchased from commercial suppliers and used without further purification. All reactions were performed in clean oven-dried round bottom flasks. NMR spectra were recorded on Bruker machines ^1^H (300 MHz, 400 MHz), ^31^P (121 MHz, 161MHz), ^13^C (100MHz). Chemical shifts are reported in ppm by using solvent residual peak as a reference (CDCl_3_: 7.26 ppm or CD_3_OD: 3.3 ppm or D_2_O: 4.79 ppm). Coupling values are reported in Hertz (Hz) (^31^P and ^13^C values reported are ^1^H decoupled). Multiplets are denoted as follows (s-singlet, br s-broad singlet, d-doublet, t-triplet, m-multiplet, p-pentet, dd-doublet of doublets). High resolution mass spectrometry data was acquired on Waters Xevo G2-S Q-TOF mass spectrometer in ESI positive or negative mode.

Chemical structures and number associated to each chemical are presented in **Tables S1** and **S2**. Generalized names were used for annotation of all the compounds.

Compounds: **1, 2, 3, 5, 6, 7, 8, 13, 14, 15, 17, 18, 20, 22, 23, 24, 25, 30, 31, 34, 35, 36, 37 and 38** are purchased from Sigma-Aldrich.

*In situ* preparation of the nerve agents (**GD (32) and GF (33)**) and synthesis of ***O-*cyclohexyl *O*-*(*3-cyano-4methyl-7-coumarinyl) methyl phosphonate (*O-*cycloxehyl *O-*coumarin methyl phosphonate,29)** was done as described previously *(1)*. Details regarding nerve agents *in situ* preparation in diluted aqueous solutions can be obtained upon request from the corresponding author.

**Synthesis of *O,O*-diphenyl phosphate (diphenyl phosphate, 4):** To a solution of diphenyl phosphorylchloride (1 ml, 4.8 mmol) in 10 ml of acetonitrile: water (1:1) mixture, NaOH (386 mg, 9.6 mmol) was added and stirred for 5min. Acetonitrile was evaporated and the reaction mixture was then acidified to a pH of ~3 by using a solution of 1M HCl . The obtained crude aqueous mixture was extracted twice with dichloromethane and the organic layer was dried over anhydrous Na_2_SO_4_. Solvents were evaporated to dryness under vacuum to obtain diphenyl phosphate as a white solid. Yield (970 mg, 81%). ^1^H NMR (300 MHz, CDCl_3_) δ 7.65 (br s, 1H), 7.38 – 7.25 (m, 4H, merged solvent signal), 7.24 – 7.11 (m, 6H). ^31^P NMR (121 MHz, CDCl_3_) δ -8.78. (HRMS, ESI -ve) m/z: [M - H]^-^ Calcd for C_12_H_10_O_4_P 249.0317; Found 249.0318.

**General procedure for the synthesis of phosphotriesters (9, 10, 11, 12, 26):** To a solution of dialkyl or diaryl phosphorylchloride (1.0 – 1.5 eq) and alcohol derivative (1.0 eq) in dichloromethane (up to 5 ml per mmol of alcohol derivative), triethylamine (1.0 - 2.0 eq) was added and stirred overnight at room temperature. Upon completion (disappearance of starting material by TLC), the reaction mixture was extracted twice, sequentially with 1M HCl, sat. NaHCO_3_ and brine solutions. The organic layer was dried over anhydrous Na_2_SO_4,_ and solvents were evaporated under vacuum. The crude product was then purified by silica gel column chromatography using Ethylacetate and Hexane gradients.

***O,O*-Dimethyl *O*-phenyl phosphate (Dimethyl phenyl phosphate, 9):** Synthesized from dimethyl phosphorylchloride (3.43 ml, 31.87 mmol),phenol (2.0 g, 21.2 mmol) and triethylamine (5.92 ml, 42.5 mmol) using general procedure, obtained as colorless oil.^1^H NMR (300 MHz, CDCl_3_) δ 7.41 – 7.29 (m, 2H), 7.25 – 7.13 (m, 3H), 3.86 (d, *J* = 11.3 Hz, 6H). ^31^P NMR (121 MHz, CDCl_3_) δ -2.77. (HRMS, ESI +ve) m/z: [M + Na]^+^ Calcd for C_8_H_11_O_4_PNa 225.0293; Found 225.0294.

***O*-(4-acetylphenyl) *O,O*-dimethyl phosphate (Dimethyl acetophenone phosphate, 10):** Synthesized from dimethyl phosphorylchloride (2.4 ml, 22.03 mmol), 4-hydroxyacetophenone (2.0 g, 14.69 mmol) and triethylamine ( 4.1 ml, 29.38 mmol) using general procedure, obtained as colorless oil. ^1^H NMR (300 MHz, CDCl_3_CDCl_3_) δ 7.96 (d, *J* = 8.5 Hz, 2H), 7.29 (d, *J* = 8.4 Hz, 2H), 3.88 (d, *J* = 11.4 Hz, 6H), 2.58 (s, 3H). ^31^P NMR (121 MHz, CDCl_3_) δ -3.32. (HRMS, ESI +ve) m/z: [M + Na]^+^ Calcd for C_10_H_13_O_5_PNa 267.0398; Found 267.0399.

***O*-(4-acetylphenyl) *O,O*-diethyl phosphate (Diethyl acetophenone phosphate, 11):** Synthesized from diethyl phosphorylchloride (3.2 ml, 22.03 mmol), 4-hydroxyacetophenone (2.0 g, 14.69 mmol) and triethylamine (4.1 ml, 29.38 mmol) using general procedure, obtained as colorless oil. ^1^H NMR (300 MHz, CDCl_3_) δ 7.96 (d, *J* = 8.6 Hz, 2H), 7.30 (d, *J* = 8.6 Hz, 2H), 4.23 (p, *J* = 7.4 Hz, 4H), 2.58 (s, 3H), 1.35 (t, *J* = 7.0 Hz, 6H). ^31^P NMR (121 MHz, CDCl_3_) δ -5.55. (HRMS, ESI +ve) m/z: [M + Na]^+^ Calcd for C_12_H_17_O_5_PNa 295.0711; Found 295.0711.

***O*-(4-acetylphenyl) *O,O*-diphenyl phosphate (Diphenyl acetophenone phosphate, 12):** Synthesized from diphenyl phosphorylchloride (3.2 ml, 15.4 mmol), 4-hydroxyacetophenone and triethylamine (4.3 ml, 30.8 mmol) using general procedure, obtained as white solid. ^1^H NMR (300 MHz, CDCl_3_) δ 7.99 (d, *J* = 8.4 Hz, 2H), 7.45 – 7.31 (m, 6H), 7.31 – 7.20 (m, 6H, merged solvent signal), 2.61 (s, 3H). ^31^P NMR (121 MHz, CDCl_3_) δ -16.91. (HRMS, ESI +ve) m/z: [M + Na]^+^ Calcd for C_20_H_17_O_5_PNa 391.0711; Found 391.0705.

***O*-methyl methyl phosphonate (16):** To a solution of dimethyl methylphosphonate (0.87 mL, 8 mmol) in methanol (4.5 mL), 5.71 mL of 2N NaOH was added dropwise. The resulting reaction mixture was stirred at room temperature for 6 h. Afterwards, the mixture was diluted with 14 mL of water and then methanol was evaporated under vacuum. The resulting aqueous phase was twice extracted with dichloromethane (2 x 3 mL) and then it was acidified with HCl until pH~2, evaporated under vacuum, providing a white solid. The residue was suspended in ethanol (3 x 3mL) and the salts were collected by filtration while the pooled organic phases were evaporated under vacuum, providing the pure product as a waxy white solid (800 mg, 7.28 mmol; 91%). ^1^H NMR (400 MHz, CD_3_OD) 3.57 (d, *J* = 10.7 Hz, 3H), 1.25 (d, *J* = 16.6Hz, 3H), ^31^P NMR (161 MHz, CD_3_OD) 26.44, ^13^C NMR (100 MHz, CD_3_OD) 50.18 (m), 10.04 (dd, *J* = 138.4, 3.1 Hz).

***O*-(4-acetylphenyl) *O*-methyl methyl phosphonate (*O*-methyl *O*-acetophenone methyl phosphonate, 19):** Oxalyl chloride (0.172 mL, 2.0 mmol) was added to a solution of dimethyl methylphosphonate (0.217 mL, 2 mmol) in dichloromethane (2 mL). The mixture was stirred overnight at room temperature. Afterwards, the mixture was concentrated under vacuum. The resulting residue was then diluted with 3 mL of dichloromethane and added dropwise to a solution of 4-hydroxyacetophenone (181.0 mg, 1.33 mmol) and triethylamine (0.371 mL, 2.67 mmol) in dichloromethane (5 mL). The resulting reaction was stirred at room temperature until TLC indicated the disappearance of starting material. The solution was diluted with dichloromethane (10 mL) and then sequentially washed with 10% aqueous HCl (3 mL), 10% aqueous NaOH (3 mL) and brine (3 mL). The organic phase was dried over anhydrous sodium sulfate and the solvent was evaporated under vacuum*,* resulting in 163.0 mg (0.71 mmol, 53.38%) of the pure product as a yellow oil. ^1^H NMR (400 MHz, CDCl_3_)7.91 (d, *J* = 8.5 Hz, 2H), 7.24 (d, *J* = 8.5 Hz, 2H), 3.76 (d, *J* = 11.4 Hz , 3H), 2.52 (s, 3H), 1.61 (d, *J* = 17.7 Hz, 3H), ^31^P NMR (161 MHz, CDCl_3_) 29.00, ^13^C NMR (100 MHz, CDCl_3_) 196.62, 154.27, 154.16, 130.89, 130.65, 130.38, 120.33, 52.82 (m), 26.47, 10.89 (dd, *J*= 145.2, 2.3 Hz).

***O*-methyl hydrogen phosphonate (Methyl phosphite, 21):** Under nitrogen atmosphere, at 0 °C, methanol (1.44 mL, 43.68 mmol) in dichloromethane (0.5 mL) was added dropwise to a stirring solution of phosphorous trichloride (1.27 mL; 14.56 mmol) in dichloromethane (0.5 mL). The reaction mixture was stirred for 10 minutes at the same temperature. Afterwards, dichloromethane was removed under vacuum and then 13.5 mL of 20% aqueous ammonia was added. The resulting solution was then stirred for additional 10 minutes and concentrated under vacuum obtaining a white solid. The crude product was triturated with ethyl ether (3 x 20 mL) to obtain the pure ammonium salt as a white solid (1.04 g, 9.19 mmol; 63.11%). ^1^H NMR (400 MHz, D_2_O) 6.52 (d, J = 634.4 Hz, 1H), 3.40 (d, J = 12.2 Hz, 3H), ^31^P NMR (161 MHz, D_2_O) 8.38 (d, J = 13.5 Hz), ^13^C NMR (100 MHz, D_2_O) 50.77 (dd, J = 18.5, 3.9 Hz).

***O,O*-diphenyl *O*-(4-nitrophenyl) phosphate (Diphenyl p-nitrophenyl phosphate or diphenyl paraoxon, 26):** Synthesized from diphenyl phosphorylchloride (3.0 ml, 14.52 mmol), 4-nitrophenol (2.0 g, 14.38 mmol) and triethylamine (2.2 ml, 15.81 mmol) using general procedure, obtained as white solid. ^1^H NMR (300 MHz, CDCl_3_) δ 8.28 (d, *J* = 8.9 Hz, 2H), 7.47 – 7.34 (m, 6H), 7.33 – 7.23 (m, 6H, merged solvent signal). ^31^P NMR (121 MHz, CDCl_3_) δ -17.14. (HRMS, ESI +ve) m/z: [M + Na]^+^ Calcd for C_18_H_14_NO_6_PNa 394.0456; Found 394.0450.

***O,O*-dimethyl *O*-*(*3-cyano-4methyl-7-coumarinyl) phosphate (Dimethyl coumarin phosphate, 27):** To a cooled solution of 3-cyano-7-hydroxy-4-methylcoumarin (200 mg, 1mmol) and triethylamine (120 mg, 1.2 mmol) in 20 ml of dichloromethane, dimethyl phosphorylchloride (144 mg, 1mmol) was added dropwise under stirring. After 24 hr at room temperature, 50 ml of diethyl ether was added, and the solid was removed by filtration, the organic layer was washed with 3% NaHCO_3_ (30 ml), water (2x30 ml) and brine (30 ml). After drying on anhydrous Na_2_SO_4_, the organic solvent was removed under vacuum leaving 250 mg of bright yellow crystals. Silica gel chromatography (5% methanol: chloroform) gave the pure compound. ^1^H NMR (300 MHz, CDCl_3_) δ 7.72 (d, *J* = 8.9 Hz, 1H), 7.35 (dd, *J* = 8.8, 2.2 Hz, 1H), 7.26 (s, 1H, merged solvent signal), 3.91 (d, *J* = 11.5 Hz, 6H), 2.76 (s, 3H). ^31^P NMR (121 MHz, CDCl_3_) δ -3.63.

***O,O*-diethyl *O*-*(*3-cyano-4methyl-7-coumarinyl) phosphate (Diethyl coumarin phosphate, 28)** was synthesized using 3-cyano-7-hydroxy-4-methylcoumarin and diethyl phosphorylchloride according to the same protocol as dimethyl coumarin phosphate (27). ^1^H NMR (300 MHz, CDCl_3_) δ 7.72 (d, *J* = 8.9 Hz, 1H), 7.35 (dd, *J* = 8.8, 2.2 Hz, 1H), 7.26 (s, 1H, merged solvent signal), 4.26 (p, *J* = 7.2 Hz, 4H), 2.75 (s, 3H), 1.38 (t, *J* = 7.0 Hz, 6H). ^31^P NMR (121 MHz, CDCl_3_) δ -5.89

**Synthesis of thiobutyl butyrolactone (35):** synthesized according to previously reported protocol *(2)*.

**Synthesis of *O*-(4-aminobutyl) *O,O*-diphenyl phosphate (ligand for CNBr-activated sepharose, 39):**

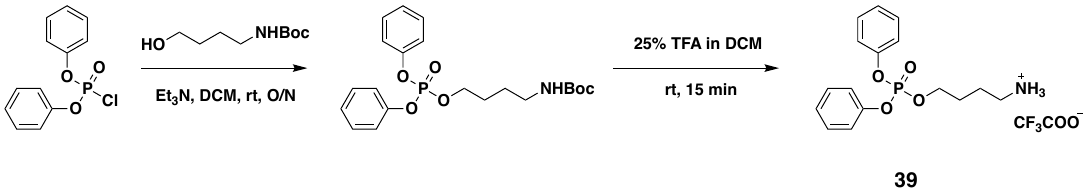
To a solution of diphenyl phosphorylchloride (110 µl, 142mg, 0.53 mmol) and 4-(Boc-amino)-1-butanol linker (100 mg, 0.53 mmol ) in dichloromethane (2 ml), triethylamine (74 µl, 53.6 mg, 0.53 mmol) was added and stirred overnight. The reaction mixture was extracted twice with brine and organic layer was dried under anhydrous Na_2_SO_4_. The solvents were evaporated to dryness under vacuum and the crude product was resuspended in a solution of 25% trifluoroacetic acid (TFA) in dichloromethane and stirred for 15 min. Excess solvents and reagents were evaporated to dryness, re-suspended in dichloromethane and re-evaporated (x3 times) to remove traces of TFA. The obtained TFA salt of 4-aminobutyl diphenyl phosphate was vacuum dried and used as is without any further purification (Impurities are likely, TFA salts of triethylamine and linker). ^1^H NMR (300 MHz, D_2_O) δ 7.43 (t, *J* = 7.8 Hz, 4H), 7.36 – 7.14 (m, 6H), 4.36 (q, *J* = 6.1 Hz, 2H), 2.93 (t, *J* = 7.2 Hz, 2H), 1.84 – 1.62 (m, 4H). ^31^P NMR (121 MHz, D_2_O) δ -9.28. (ESI +ve) m/z: [M + H]^+^ Calcd for C_16_H_21_NO_4_P 322.12; Found 322.34.

**Isolation of OP-degrading bacteria from the marine environment**

We sampled seawater from the Red Sea at the natural reserve in Eilat, Israel (pier UIU at 4 m depth, on February 3^rd^, 2021, <http://www.meteo-tech.co.il/eilat-yam/eilat_en.asp>, N 29.5035° E 34.9178°), and three locations at the Achziv beach on February 10^th^, 2021 (N 33.0438°, E 35.1011°) at the Mediterranean Sea – MS sample was taken 600 m from the shore (density 1.0278 kg/L, T 22°C, EC 57.5 mS/cm, pH 6.48, DO 10.42 mg/L) and two samples were acquired at locations of lower salinity in which mixing of fresh water and seawater occurs (MSM1 - 0cm depth, ~10 m offshore, density 1.0106 kg/L, T 21.7°C, EC 25.6 mS/cm, pH 7.35, Do, 6.27 mg/L; MSM2 - 18 cm depth, ~10 m offshore, density 1.0193 kg/L, T 19.7°C, EC 37.1 mS/cm, pH 7.45, DO 3.22 mg/L). Water samples were filtered through 40 µm filters and diluted 100-fold in 10 ml ASW (as described above, 30 mM HEPES instead of K_2_HPO4 was used for basal medium) supplemented with 1 mM OPs as the sole P source (mix trimethyl-/triethyl phosphate (compounds 8 and 36), dimethyl phenyl phosphate (compound 9), dimethyl acetophenone phosphate (compound 10) and diphenyl acetophenone phosphate (compound 12)) and in combination with 3 different carbon sources (10 mM glucose, succinate or glycerol). The cultures were grown at 30°C with continuous shaking (220 rpm) and when they reached OD_600_ 0.1-0.5, they were diluted into the same fresh medium. After 4 serial passages, cultures were streaked on marine broth agar plates and grown at 30°C. Three colonies were picked from each plate for 16S rRNA amplification with primer set B_27_F 5’-AGAGTTTGATCCTGGCTCAG-3’ and U_1492_R 5’-GGTTACCTTGTTACGACTT-3’. The same three colonies were re-inoculated into the same medium used for their enrichment (containing the same phosphorus and carbon sources). Isolates were identified by 16S rRNA sequencing, **Table S3** and stored in glycerol stocks. For further verification, isolated colonies were again plated on marine broth agar plates and three new colonies were collected for 16S rRNA verification and stored in glycerol stock.

**Growth profiling of the model marine bacteria and *E. coli* BL21 on various organophosphates (OPs) as the phosphorus source**

Marine strains *Ruegeria* sp. TM1040 and *Ruegeria pomeroyi* DSS-3 were kindly received from Jonathan Todd and Andrew Johnston (University of East Anglia, Norwich Research Park, Norwich, UK). The bacterial strains of *Phaeobacter inhibens* DSM 17395 and *Dinoroseobacter shibae* DSM 16493T were purchased from the German collection of microorganisms and cell cultures (DSMZ, Braunschweig, Germany).

Strains were validated by sequencing 16s rRNA, using the primer set- B_27_F 5’-AGAGTTTGATCCTGGCTCAG-3’ and U_1492_R 5’-GGTTACCTTGTTACGACTT-3’. Bacteria from glycerol stocks were cultured on Marine Broth (Difco) agar plates overnight at 30°C on. Colonies from the agar plates were inoculated into Marine Broth and incubated at 30°C overnight with continuous shaking (200 rpm). Overnight cultures were washed twice with artificial seawater, OD was adjusted to 1 and then diluted 100-fold in fresh artificial seawater medium (ASW) supplemented with 10 mM CaCl_2_, basal media, bacterial vitamin mix *(3)*, trace metal solution *(4)*, 500 µM NaNO_3_ and 10 mM carbon source: glucose (*P. inhibens*), succinate (*R. pomeroyi* DSS3 and *D. shibae*), glycerol (*Ruegeria* sp. TM1040) and 1 mM OPs (substrates are listed in **Table S1,** compounds 1-23) as the phosphorous source. Growth monitoring of *Ruegeria* sp. TM1040, *Ruegeria pomeroyi* DSS-3 and *Phaeobacter inhibens* was conducted using 10 ml cultures in 50 ml Falcon tubes cultured for 78 h at 30°C with continuous shaking (200 rpm). OD was measured in intervals of 4 to 8 h. Two biological replicates were measured for each sample and mean values with standard deviations (SD) are presented. *Dinoroseobacter shibae* OD was monitored in a 96-well microtiter plate reader for 78 h at 30°C and measured every 1 hour. Each well contained 150 µl culture covered with 50 µl hexadecane to avoid evaporation during incubation. Biological triplicates were measured for each sample and mean values with SD are presented.

Profiling of *E. coli* BL21 was carried out with similar protocols used for marine bacteria but using media suitable for this strain. Glycerol stocks were cultivated on LB plates, and colonies were inoculated into LB medium and grown. Growth monitoring was conducted using MOPS medium supplemented with 0.4 % glucose and 1 mM OPs. The OD was monitored in a 96-well microtiter plate reader with biological triplicates for each sample.

The maximal OD for each condition was used to generate the heat map in **Fig. 3**.

**Growth profiling of the isolated marine bacteria on various organophosphates (OPs) as the phosphorus source**

The protocol described above was used. All cultures were cultivated in 96-well microtiter plates (each well contained 150 µl culture covered with 50 µl hexadecane to avoid evaporation). OD_600_ was monitored in a plate reader (Epoch™ 2 Microplate Spectrophotometer, BioTek) every hour for 49-67 h with continuous shaking at 30°C. Strains were monitored using technical triplicates (including *Celeribacter naphthalenivorans*, *Alteromonas macleodii*, *Alteromonas mediterranea*, *Vibrio sp.*, *Labrenzia* *sp.*. To validate similar growth dynamics among biological replicates, a subset of strains was monitored using biological triplicates (including *Cobetia* sp, *Marinobacter* *sp.*, *Pseudoalteromonas* sp., *Marinomonas brasilensis*, *Tateyamaria omphalii*). Importantly, similar growth dynamics were observed among technical and biological replicates. The growth curves showed the same trend as in the case of technical triplicates. Mean values with SD are presented in the growth curves. The max OD_600_ values were used to generate heat maps. Particular wells showed appearance of a second peak of growth. Aggregates were evident in these wells thus the second peak likely appeared due to precipitation in the microtiter plate. In these cases, values for the heat map in **Fig. 3** were taken from the first peak, before the appearance of the second peak (examples are indicated by an arrow in **Fig. S6**).

**Activity profiling of the *Ruegeria sp.* TM1040 and *R. pomeroyi* DSS3 cell lysate**

A single colony was inoculated into Marine broth for overnight at 30°C with continuous shaking (220 rpm). The culture was then diluted 100-fold in ASW supplemented with 10 mM succinate or glycerol (*R. pomeroyi* DSS3/*Ruegeria sp.* TM1040) and 1 mM K_2_HPO_4_. Cells were grown at 30°C until OD_600_ ~ 1. Cells were pelleted at 2500×g for 30 minutes at 4°C and the pellets were frozen at -20°C. Frozen cells originating from 250 ml culture were thawed and resuspended in 20 ml of lysis buffer (50 mM Tris pH 8, 50 mM NaCl), supplemented with Protease Inhibitor Cocktail (EDTA-free; ABCAM, diluted 1:1000), benzonase (MERCK, 100 units) and lysozyme (Glentham Life Sciences Ltd, 0.4/1 mg/ml). Resuspended cells were incubated for 1 hour at 37°C and lysis was completed by sonication (15 sec × 8 cycles, 35% amplitude, on ice). Cell debris were removed by centrifugation (45 minutes at 7500× g at 4°C) and the supernatant was used to measure activity.

Lysate activity was measured in the microtiter plate reader (Epoch™ 2 Microplate Spectrophotometer, BioTek) over 30 minutes, 90 µl substrate was added to each well containing 10 µl supernatant. We used 6 substrates for activity profiling of the lysate: methyl paraoxon (compound 24), paraoxon (compound 25), phenyl paraoxon (compound 26), dimethyl coumarin phosphate (compound 27), diethyl coumarin phosphate (compound 28) and *O-*cyclohexyl *O-*coumarin methyl phosphonate (compound 29). Substrates were prepared in final concertation of 1 mM in activity buffer (50 mM Tris pH 8, 50 mM NaCl), with the exception of phenyl paraoxon, which was prepared in final concentration of 0.1 mM in activity buffer with 10% DMSO, due to its solubility. The release of p-nitrophenol (at 405 nm; extinction coefficient 16000 M^-1^cm^-1^) and coumarin (at 400 nm; extinction coefficient 37000 M^-1^cm^-1^) was monitored. All measurements were conducted in triplicates and values were normalized according to the substrate that was best utilized whose activity was set to 1.

**Isolation of phosphotriesterases from *R. pomeroyi* DSS3 and *Ruegeria sp.* TM1040**

Cell pellets were prepared as described in the previous section.

Enzymatic fractionation and purification based on specific activity from *Ruegeria* *sp.* TM1040: The cell pellet from 250 ml culture was resuspended in 10 ml of lysis buffer (100 mM Tris pH 8, 50 mM NaCl, 1mM CaCl_2_, 1mM MgCl_2_, 0.2 mg/ml lysozyme, 10 µl Protease Inhibitor Cocktail, 1 µl benzonase), sonicated (3 min: 30 sec on, 30 sec off) and incubated for 2h at 37°C with continuous shaking. Cell debris were removed by centrifugation (45 min at 7500 x g at 4°C) and the supernatant was filtered through a 0.22 µm syringe filter.

The supernatant was subject to a 2-step purification. *In the 1^st^ step*, the filtrate was loaded on an anion exchange HiTrap Q HP 5 ml column (GE Healthcare Life Sciences, Boston, Massachusetts), at a flow rate of 5 ml/min, using an FPLC AKTA-prime plus device (GE, Boston, Massachusetts) pre-equilibrated with buffer A_Q_ (100 mM Tris pH 8, 1mM CaCl_2_, 1mM MgCl_2_). The enzyme variants were eluted by a gradient of buffer A_Q_ supplemented with 1 M NaCl (0-100 % gradient within 20 column volumes, at flow rate 5 ml/min). The phosphotriesterase activity of the collected fractions was measured with 0.1 mM phenyl paraoxon (compound 26) in activity buffer (100 mM Tris pH 8, 1mM CaCl_2_, 1mM MgCl_2_). The increase in absorbance at 405 nm was followed using a plate-reading spectrophotometer (PowerWave HT, BioTek, Winooski, Vermont). The active fractions were combined and dialyzed in SnakeSkin Dialysis Tubing 10K MWCO (Thermo Fisher Scientific, Waltham, Massachusetts) against activity buffer. *In the 2^nd^ step*, the ligand (butylamine diphenyl phosphate) was coupled to 50 µl of CNBr-activated Sepharose 4B beads following the standard ligand coupling procedure. Sepharose beads with butylamine diphenyl phosphate (compound 39) as a ligand were equilibrated in the activity buffer and fractions with the highest activity from the previous purification step were applied on the column. After extensive (~30 column volumes) washing with activity buffer 0.5 mL of 0.1 mM phenyl paraoxon in the activity buffer was applied to the column. Characteristic yellow color of p-nitrophenol release upon incubation with sepharose indicated binding of the enzyme to the resin. Sepharose beads with bound enzyme (50 µl) were mixed with 25 µl of activity buffer and 25 µl of sample buffer (4X) for SDS-PAGE. The samples were boiled, centrifuged and 20 µL of the supernatant was tested on SDS-PAGE. The 35 kDa band was cut from the gel and submitted for proteomics analysis. As controls for the background proteins, the bands with the similar sizes were cut from the gel from the inactive and active HP HiTrap column fractions.

Enzymatic fractionation and purification based on specific activity from *R. pomeroyi* DSS3: The cell pellet from 2 l culture was resuspended in 80 ml of lysis buffer (50 mM Tris pH 8, 1 mM MgCl_2_, 1 mM CaCl_2_, 0.4 mg/ml lysozyme, 10 µl Protease Inhibitor Cocktail, 1 µl benzonase), sonicated (5 min: 15 sec on, 45 sec off, 35% amplitude) and incubated for 2h at 37°C with continuous shaking. The cell debris was removed by centrifugation (45 min at 7500 x g at 4°C) and the supernatant was filtered through a 0.45 µm syringe filter.

A three-step protocol was employed to isolate an unknown phosphotriesterase from the cell lysate of *R. pomeroyi* DSS3. *First* we optimized heat precipitation – the cell lysate was incubated at various temperatures for 30 min in the heating block (25, 70, 75, 80 and 85°C), cooled on ice and measured residual phosphotriesterase activity with 1 mM methyl paraoxon (compound 24), **Figure S24**. Unknown phosphotriesterase was activated upon heating while other proteins precipitated, which led to partial purification of the enzyme. Heating at 80°C gave the best ratio of activation and purity. Therefore, *in the 1^st^ step*, cell lysate was heated at 80°C for 30 min, precipitated proteins removed by centrifugation (13 000xg, 15 min, 4°C) and the supernatant was filtered through a 0.45 µm syringe filter. *In the 2^nd^ step*, the filtrate was loaded on an anion exchange HiTrap Q HP 5 ml column (GE Healthcare Life Sciences, Boston, Massachusetts), at a flow rate of 2 ml/min, using an FPLC AKTA-prime device (GE, Boston, Massachusetts) pre-equilibrated with buffer A_Q_ (50 mM Tris pH 8, 1mM CaCl_2_, 1mM MgCl_2,_). The enzyme variants were eluted by a gradient of buffer A_Q_ supplemented with 1 M NaCl (0-100 % gradient within 20 column volumes, at flow rate 5 ml/min). The phosphotriesterase activity of the collected fractions was measured with 1 mM methyl paraoxon (compound 24) in activity buffer (50 mM Tris pH 8, 50 mM NaCl). The increase in absorbance at 405 nm was followed using a plate-reading spectrophotometer (PowerWave HT, BioTek, Winooski, Vermont). The active fractions were analyzed by SDS-PAGE, combined and dialyzed in SnakeSkin Dialysis Tubing 10K MWCO (Thermo Fisher Scientific, Waltham, Massachusetts) against A_B_ (50 mM Na_2_HPO_4_, pH 7, 1 M ammonium sulfate, 1 mM CaCl_2_, 1 mM MgCl_2_) *In the 3^rd^ step*, the sample was loaded on hydrophobic interaction column HiTrap Butyl-HP 5 ml (GE Healthcare Life Sciences, Boston, Massachusetts). After washing with 5 column volumes, the enzyme was eluted by gradient decrease of ammonium sulfate in 10 column volumes. Phosphotriesterase activity was monitored for each fraction and active fractions were analyzed by SDS. However, due to low protein concentration fractions with the highest activity were concentrated 14-16-fold and again analyzed by SDS-PAGE and *in situ* gel activity.

*In situ* activity on SDS-PAGE gel: Active concentrated fractions after hydrophobic butyl chromatography were mixed with SDS-dye without boiling, and were loaded on 12% SDS acrylamide gel in 2 replicates. After separation, the gel was washed in 3 times with 20 ml renaturation buffer (0.2 mM Bicine, 0.2% Triton X-100, 100 mM NaCl, 1 mM MgCl_2_, 1 mM CaCl_2_, pH 8). The renatured gel was divided in two parts. The first half of the gel was stained with Commassie blue, and the second half was placed on 1.5% agar gel with 100 mM NaCl pH 8, 0.4 mg/ml Cresol purple, 1 mM methyl-paraoxon (compound 24). Additionally, acrylamide gel was covered with few ml of 100 mM NaCl and 1 mM methyl-paraoxon from the top. After few minutes yellow color was developed slightly above 25 kDa protein marker. Yellow p-nitrophenol was released during the reaction and also proton was generated that led to decrease in pH and change of cresol from dark red to yellow color, intensifying the yellow bands appearing on both the acrylamide and the agarose gel.

For analysis by mass spectrometry, fractions after 3^rd^ step of purification were sent. The samples were cut from the gel (Commassie and *in situ* activity stained).

**Identification of phosphotriesterases by mass spectrometry**

Sample preparation: Gel bands were excised from the gel, sliced into 1-2 mm pieces and placed in a microcentrifuge tube. Gel bands were destained with 25mM NH_4_HCO_3_ in 50% acetonitrile (ACN) and then vacuum dried. Protein disulfide bonds were reduced by saturating the dry gel bands with 10 mM dithiothreitol in 25 mM NH_4_HCO_3_ at 56°C for 1 h and alkylated with 55 mM iodoacetamide in 25 mM NH_4_HCO_3_ in the dark for 45 min at room temperature. Bands were washed twice with 25 mM NH_4_HCO_3_ and twice with 25 mM NH_4_HCO_3_ in 50% ACN. Bands were vacuum dried and rehydrated with 500 ng trypsin (Promega; Madison, WI, USA) in 25 mM NH_4_HCO_3_ at 4°C for 10 min followed by overnight incubation at 37°C. 500 ng trypsin was added for a second digestion for 4 h at 37°C. Peptides were then extracted by addition of 50% ACN/5% formic acid, vortexed, centrifuged, and the supernatant collected. The digestions were stopped by addition of trifluroacetic acid (1% final concentration). The samples were stored in -80˚C until further analysis.

Liquid chromatography: ULC/MS grade solvents were used for all chromatographic steps. Dry digested samples were dissolved in 97:3% H_2_O/acetonitrile + 0.1% formic acid. Each sample was loaded and analyzed using split-less nano-Ultra Performance Liquid Chromatography (10 kpsi nanoAcquity; Waters, Milford, MA, USA). The mobile phase was: A) H_2_O + 0.1% formic acid and B) acetonitrile + 0.1% formic acid. Desalting of the samples was performed online using a Symmetry C18 reversed-phase trapping column (180 µm internal diameter, 20 mm length, 5 µm particle size; Waters). The peptides were then separated using a T3 HSS nano-column (75 µm internal diameter, 250 mm length, 1.8 µm particle size; Waters) at 0.35 µL/min. Peptides were eluted from the column into the mass spectrometer using the following gradient: 4% to **3**3% B in 55 min, **3**3% to 90% B in 5 min, maintained at 90% for 5 min and then back to initial conditions.

Mass Spectrometry: The nanoUPLC was coupled online through a nanoESI emitter (10 μm tip; New Objective; Woburn, MA, USA) to a quadrupole orbitrap mass spectrometer (Q Exactive HF, Thermo Scientific) using a FlexIon nanospray apparatus (Proxeon).

Data was acquired in data dependent acquisition (DDA) mode, using a Top10 method. MS1 resolution was set to 120,000 (at 200m/z), mass range of 375-1650m/z, AGC of 3e6 and maximum injection time was set to 60 msec. MS2 resolution was set to 15,000, quadrupole isolation 1.7m/z, AGC of 1e5, dynamic exclusion of 20 sec and maximum injection time of 60 msec.

Data processing and analysis: The data was processed using Proteome Discoverer version 2.4.1.15 and searched using SequestHT and MS-Amanda against a protein database containing the respective analyzed bacterium as downloaded from Uniprot.org (*Ruegeria pomeroyi* DSS3 and *Ruegeria* sp. *TM1040*), appended with a list of common lab contaminants. Enzyme specificity was set to trypsin and up to two missed cleavages were allowed. Fixed modification was set to carbamidomethylation of cysteines and variable modifications were set to oxidation of methionines, and deamidation of asparagines and glutamine. Peptide and protein identifications were filtered at an FDR of 1% using Percolator.

**Cloning of MBL-PTE and cyclase-PTE in *E. coli* expression vector**

The metallo-beta-lactamase gene identified by MS-proteomics (gene accession number: TM1040_1477; UniProt ID: Q1GGK6) was amplified from *Ruegeria* sp. TM1040 genome without signal sequence (F 5'-ggagatatacatatggcaccaatggcccctgcgcctgt-3’ and R 5'-gtggtggtgctcgagtcagagg-ccgaattgccagggcg-3’). Addition of C terminal 6xHis tag was performed on the new construct using outward complimentary primers (MEGAWHOP) (F 5’-ggaggaggacatcat-catcatcatcattgactcgagcaccaccaccaccaccact-3’ and R 5’-tcaatgatgatgatgatgatgtcctcctccgaggccg-aattgccagggcgc-3’).

The cyclase-PTE gene identified by MS-proteomics (gene accession number: SPO0761; UniProt ID: Q5LVE1) was amplified from *Ruegeria pomeroyi* DSS3 genome with the primers (F 5’-tatacatatggccggcattggcgaagtgcg-3’ and R 5’-atatctcgagtcagaccatggcaaagatgcggg-3’) without signal peptide sequence.

PCR products were digested with NdeI and XhoI restriction enzymes (fast digest; Fermentas) and cloned to pET21a. The recombinant plasmids were transformed into electro-competent *E. coli* BL21 (DE3) and cultured in LB medium with 100 µg/ml ampicillin.

**Catalytic metal identification**

Expression medium (LB for cyclase-PTE and 2YT for MBL-PTE) was supplemented with 0.1 mM metals, at OD600nm 0.6 protein expression was induced with 1 mM IPTG for overnight at 18/37°C (cyclase/lactamase). Cells were pelleted in lysed in 50 mM Tris, 50 mM NaCl pH 8.0 with addition of 1 mM corresponding metal. Phosphotriesterase activity with methyl paraoxon (compound 24)/diphenyl paraoxon (compound 26) (cyclase/lactamase) was measured in the cell lysate. Measurements were done in duplicate and mean values with standard error were presented.

**Recombinant expression and purification of MBL-PTE from *Ruegeria* sp. TM1040**

*E. coli* BL21 cells expressing MBL-PTE were inoculated into 5 ml of 2xYT medium supplemented with 100 µg/ml ampicillin. The culture was grown overnight at 37°C shaking at 220 rpm. The overnight culture was diluted 100-fold in 500 ml 2xYT with 1 mM MnCl_2_ and 100 µg/ml ampicillin, and grown at 37°C with continuous shaking (220 rpm). At OD_600_ of 0.6, expression of MBL-PTE was induced with 1 mM IPTG. After overnight growth at 37°C, the cultures were centrifuged at 3000x g for 30 minutes and the pellet was frozen at -20°C. The cells were thawed and lysed in 10 ml of lysis buffer (50 mM Tris, 50 mM NaCl pH 8.0 with addition of 1 mM MnCl_2_, 10% glycerol, 1 mg/ml lysozyme, protease inhibitor cocktail and bensonase. Cell debris were removed by centrifugation (45 minutes at 7500× g at 4◦C) and the supernatant was collected. All procedures were performed on ice or in the cold room with ice cold buffers.

Purification of the expressed protein was done on 4 ml Ni-NTA resin, pre-equilibrated with 50 mM Tris pH 8, 500 mM NaCl, 1 mM MnCl_2_, 20% glycerol and 5 mM imidazole. The supernatant was supplemented with 5 mM imidazole and NaCl added up to 500 mM, before loaded on the column. Washing on the column was done in the same buffer as the pre-equilibration, 20 column volumes. Protein is eluted in 50 mM Tris, 50 mM NaCl, 500 mM imidazole pH 8.0. Buffer exchange was done on PD-10 desalting column, equilibrated with 50 mM Tris pH 8.0, 50 mM NaCl, 1 mM MnCl_2_ and 20% glycerol, immediately after Ni-NTA purification. The enzyme was aliquoted on ice, frozen in the liquid nitrogen and stored at -80°C in the final protein concentration of 2.7 mg/ml.

**Recombinant expression optimization and purification of cyclase-PTE from *Ruegeria pomeroyi* DSS3**

Chemically competent BL21 *E. coli* cells carrying chaperones expressing plasmids were transformed with expression vector pET21a-cyclase-PTE. The chaperones expressing plasmids are: pGro7 (expressing genes: *groES-groEL*), pKJE7 (expressing genes: *dnaK-dnaJ-grpE*), pTf16 (expressing gene: *tig*) with chloramphenicol resistance and under *araB* promoter (0.5 mg/ml L-arabinose induction), and pG-KJE8 (expressing genes: *dnaK-dnaJ-grpE, groES-groEL*) with chloramphenicol resistance and both L-arabinose and tetracycline induction (5 µg/ml tetracycline) (Chaperone plasmid set, Cat. #3340, TAKARA). In addition, chemically competent BL21 *E. coli* cells with a control plasmid pACYC (empty plasmid) were transformed with pET21a-cyclase-PTE.

Cells co-expressing both plasmids – cyclase-PTE and chaperons- were grown at 37°C in 10 ml LB, supplemented with 100 µg/ml ampicillin, 30 µg/ml chloramphenicol, 0.5 mg/ml L-arabinose, 5 µg/ml tetracycline (both inductions were used for all plasmids to standardize conditions), 0.1 mM ZnCl_2_ and 0.1% glucose. At OD_600_ of 0.6 cells were induced with 1 mM IPTG and the cultures were split to two 5 ml subcultures– for overnight expression at 18°C and 30°C, after which cells were pelleted and frozen. Cell pellet was resuspended in 500 µl of 50 mM Tris pH 8.0, 100 mM NaCl, 0.1 mM ZnCl_2_, supplemented with Protease Inhibitor Cocktail (EDTA-free; ABCAM, diluted 1:1000), benzonase nuclease (MERCK, 100 units) and lysozyme (Glentham Life Sciences Ltd, 0.4 mg/ml). Resuspended cells were incubated for 1 h at 37°C and lysis was completed by sonication (15 sec × 2 cycles, 35% amplitude, on ice). Cell debris were removed by centrifugation (45 minutes at 7500× g at 4°C) and the supernatant was used to measure activity. Phosphotriesterase activity was measured with 1 mM methyl paraoxon (compound 24) and release of p-nitrophenol was monitored at 405 nm over 30 minutes.

Purification of cyclase-PTE: Cells co-expressing pET21a-cyclase-PTE and pGro7 plasmids were grown at 37°C in 500 ml LB, supplemented with 100 µg/ml ampicillin, 30 µg/ml chloramphenicol, 0.5 mg/ml L-arabinose, 0.1% glucose and 0.1 mM ZnCl_2_. At OD_600 nm_ 0.6 cells were induced with 1 mM IPTG and the cultures was grown at 18°C overnight. Cells were pelleted at 7500xg for 30 min at 4°C and frozen at -20°C. The cell pellet was resuspended in 40 ml lysis buffer contained 50 mM Tris pH 8 and 0.1 mM ZnCl_2_ supplemented with Protease Inhibitor Cocktail (EDTA-free; ABCAM, diluted 1:1000), benzonase nuclease (MERCK, 100 units) and lysozyme (Glentham Life Sciences Ltd, 0.4 mg/ml). Resuspended cells were incubated for 1 h at 37°C and lysis was completed by sonication (15 sec × 10 cycles, 35% amplitude, on ice). Cell debris was removed by centrifugation (45 minutes at 7500× g at 4°C) and the supernatant was divided to 1.5 ml Eppendorf tubes for heating at 80°C for 30 minutes in the heating block (Thermo shaker, mrc, Burnt Mill, Elizabeth Way Harlow, Essex, UK). The lysate was then clarified by centrifugation at 13,000xg for 10 min at 4°C and the clear lysate was filtered through a 0.45 µm syringe filter (Millex-HV Syringe Filter Unit, 0.45 µm, PVDF, 33 mm, Merck KGaA, Darmstadt, Germany) and loaded on HiTrap Q-HP, 5 ml column (GE) anion exchange column pre-equilibrated with buffer A_Q_- 50 mM Tris, pH 8, 0.1 mM ZnCl_2_, at 2 ml/min. The loaded column was washed with 5 column volumes (cv) of buffer A_Q_, then the protein was eluted by a gradient of increasing NaCl concentration up to 1 M for 20 cv. Activity of each fraction was measured with 1 mM methyl paraoxon (compound 24). The most active fractions were eluted at 35 % NaCl, and analyzed by SDS Page. Protein concentration and activity was measured for each fraction and was compared to load. The active fractions were combined and dialyzed overnight in SnakeSkin Dialysis Tubing 10K MWCO against buffer A_Butyl_ 50 mM Na_2_HPO_4_, pH 7, 1 M ammonium sulfate, 0.01 mM ZnCl_2_, 1 mM MgCl_2_ and 1 mM ATP (MgCl_2_ and ATP were added in order to remove chaperones bound to cyclase-PTE). The sample was loaded at 2 ml/min on hydrophobic interaction column (HiTrap Butyl-HP, 5 ml) pre-equilibrated with buffer A_Butyl_ and the protein was eluted by gradient decrease of ammonium sulfate to 10 mM. Protein concentration and activity was measured with 1 mM methyl paraoxon (compound 24) for each fraction and was compared to load. The most active fractions were eluted at lowest concentration of ammonium sulfate. Pure and active samples were combined and dialyzed overnight in SnakeSkin Dialysis Tubing 10K MWCO against 50 mM Tris, pH 7.5, 50 mM NaCl, 0.1 mM ZnCl_2_. Active fractions are analyzed by SDS-PAGE, and fractions with the highest activity are pulled together, concentrated and protein stored at 4°C.

**Kinetic characterization of phosphotriesterases – cyclase-PTE and metallo-beta-lactamase PTE**

Activity buffers: cyclase-PTE: 50 mM Tris, 50 mM NaCl+0.1% Tergitol, pH 8.0; Metallo beta-lactamase PTE: 50 mM Tris, 50 NaCl, 1 mM MnCl_2_ + 10% glycerol, pH 8.0.

All measurement performed in duplicates at 25^o^C. Graphics and calculations were constructed and analyzed by GraphPad Prism 7.

Caution: Concentrations of the *in situ* generated G-and V-agents in diluted aqueous solutions are non-hazardous. Yet, due to their high potency as inhibitors of acetyl-cholinesterase (AChE), all safety requirements were strictly observed. The same practice should also be applied when working with the other OPs employed in this study.

Methyl paraoxon (compound 24) and paraoxon (compound 25) and hydrolysis: The release of p-nitrophenol during the hydrolysis of two OPs were monitored at 405 nm and the Michaelis-Menten plot obtained over 0.25-5 mM paraoxon and methyl paraoxon for MBL-PTE and over 0.025-1.5 mM paraoxon and methyl paraoxon for cyclase-PTE. Data points were analyzed by non-linear regression analysis and k_cat_ calculated using extinction coefficient of p-nitrophenol at pH=8, 1.6x10^4^ M^-1^cm^-1^. Cyclase- concentration was 1.6 nM for methyl paraoxon and 16 nM for paraoxon measurement, MBL-PTE concentration was 0.42 µM for methyl paraoxon and 0.83 µM for paraoxon.

Phenyl paraoxon (compound 26) hydrolysis: The release of p-nitrophenol during the hydrolysis of phenyl paraoxon monitored at 405 nm and the Michaelis-Menten plot obtained over 12.5-125 µM substrate, MBL-PTE concentration was 84 nM. Cyclase-PTE (15.9 µM) showed no release of p-nitrophenol with 0.2 mM substrate. Due to low solubility of the substrate 20 % DMSO was added to the reaction buffer.

Parathion (compound 30) and methyl parathion (compound 31) hydrolysis: The release of p-nitrophenol during the hydrolysis of two OPs were monitored at 405 nm. MBL-PTE and cyclase-PTE do not hydrolyze parathion; 8.3 µM MBL-PTE showed no release of p-nitrophenol with 0.2 mM parathion and 15.9 µM cyclase-PTE showed no release of p-nitrophenol with 1.5 mM parathion. Michaelis-Menten plot for hydrolysis of methyl parathion by cyclase-PTE was obtained over 0.1-1.5 mM substrate and 15.9 µM enzyme. MBL-PTE activity with methyl-parathion (initial velocity) was measured in the presence of 0.95 mM substrate and 0.84 µM enzyme.

GD (compound 32) and GF hydrolysis (compound 33): The *in situ* conversion of the coumarin surrogates of GD and GF to the corresponding G nerve agents in diluted aqueous solutions and the monitoring of the rate of their detoxification by OP hydrolases were performed as previously described by Gupta et al.*(5)* and Ashani et al.*(6)*. In these assays the residual nerve agents as function of time of incubation with the catalytic protein were measured by back titration of acetylcholinesterase with known active sites concentration.

Catalytic efficiencies (k_cat_/K_M_) for cyclase-PTE were determined by measuring the activity at several low GF (0.1, 0.5 and 1 µM) and GD (0.25 and 1 µM) concentrations in the approximated first-order kinetics region of the Michaelis-Menten equation. Thus, for example, the hydrolysis of 100 nM GF was initiated by 6.8 nM cyclase-PTE and at selected time intervals aliquots diluted 20-fold in 4 nM rhAChE in 50 mM Tris, 50 mM NaCl+0.1% Tergitol, pH 8.0. The near-by stoichiometry of the inhibition of the AChE activity was completed within 60-80 min. In the case of GD the reaction was initiated with 34 nM cyclase-PTE and at selected time intervals aliquots diluted 20-fold in 4 nM rhAChE in 50 mM Tris, 50 mM NaCl+0.1% Tergitol, pH 8.0. The near-by stoichiometry of the inhibition of the AChE activity was completed within 80-120 min. The % inhibited enzyme without the cyclase-PTE was assigned the 100% value of GF at t=0. The decrease in percent residual inhibitor were ploted vs time of incubation and analyzed in accordance with a mono-exponential decay kinetics (k_obs_), k_cat_/K_M_ =k_obs_/[enzyme].

Since MBL-PTE was by far less active in detoxification of GD (5 and 10 µM) and GF (10 µM), its concentration increased to 0.84 for GD and 1.68 µM for GF. At specified time intervals, the reaction mixture diluted 10- to 20 fold in 50 mM Tris-50 mM NaCl+0.1% Tergitol , and then followed immediately by dilution into the AChE solution to determine residual nerve agents as described above.

Isatin (compound 34) hydrolysis: The enzymatic hydrolysis of isatin to isatinate was monitored at 368 nm as described by Bjerregaard-Andersen *(7)*. Isatin concentration ranged 0.025-1.2 mM and the cyclase-PTE was at 0.16 µM. The data points were fitted to Michaelis-Menten plot and analyzed by non-linear regression. k_cat_ was calculated using extinction coefficient of the isatinate at pH=8, 4.5x10^3^ M^-1^cm^-1^. Metallo-beta-lactamase (0.55 µM) did not show activity when reacted with 0.69 mM isatin over 5 min.

TBBL hydrolysis (compound 35): The hydrolysis of TBBL was followed by use of Ellman’s reagent DTNB, as previously described by M^-1^cm^-1^ at pH=8.0. Michaelis-Menten plot for cyclase-PTE was obtained over 0 -0.14 mM substrate, enzyme concentration was 34 nM. Michaelis-Menten plot for MBL-PTE was obtained over 0-2.0 mM substrate and 0.42 µM enzyme.

VX (the S and R enantiomers, compound 40): The hydrolysis of the toxic and less toxic VX enantiomers (S and R, respectively at 0.01 to 0.02 mM ) in the presence of 0.25 µM cyclase-PTE and 0.55 µM MBL-PTE was followed by use of Ellman’s reagent DTNB, as previously described by Cherny et al.*(8)*. No release of the expected thiol leaving group could be observed at least in the first 10 min of incubation at 25^o^C.

Dimethyl acetophenone phosphate hydrolysis (compound 10): The release of the leaving group p-hydroxy acetophenone was followed at 325 nm using molar extinction coefficient of 1.52x10^4^ M^-1^cm^-1^ at pH=8.0. Michaelis-Menten plot performed at 0.2-3 mM dimethyl acetophenone phosphate, and the cyclase-PTE concentration was 0.16 µM. MBL-PTE at 0.42 µM was employed to generate the M-M curve over 0-1.72 mM dimethyl acetophenone phosphate.

**Co-expression of phosphotriesterases (cyclase-PTE and MBL-PTE) with GpdQ phosphodiesterase in *E. coli* BL21 and growth on OPs as the sole P source**

Cells co-expressing cyclase-PTE with GroEL-ES and cells expressing MBL-PTE were prepared as electro competent cells and each transformed separately with pET29b-GpdQ vector (phosphodiesterase from *Enterobacter aerogenes (9)*).

Cells co-expressing Cyclase-PTE, GroEL-ES and GpdQ were inoculated into 5 ml LB supplemented with 100 µg/ml ampicillin and 50 µg/ml kanamycin for overnight growth at 37°C shaking at 220 rpm. After overnight growth the cells were washed 2 times with MOPS pH 7.4 buffer supplemented with 0.4% glucose, 1 mM IPTG, 0.5 mg/ml L-arabinose, 5 µM K_2_HPO_4_. The cells were adjusted to OD_600 nm_ 1, and diluted 100-fold in 150 µl of MOPS pH 7.4 supplemented with 0.4% glucose, 1 mM IPTG, 0.5 mg/ml L-arabinose, 5 µM K_2_HPO_4_, 0.1 mM ZnCl_2_, 0.1 mM MnCl_2_ and 1 mM P source: K_2_HPO_4_, dimethyl phosphate (compound 2) and dimethyl acetophenone phosphate (compound 10). Each well was covered with 50 µl of hexadecanee (Sigma) and the 96-well microtiter plate was incubated in the plate reader at 26°C (cyclase-PTE) or 30°C (MBL-PTE), for 70 hours with continuous shaking.

**Figures**

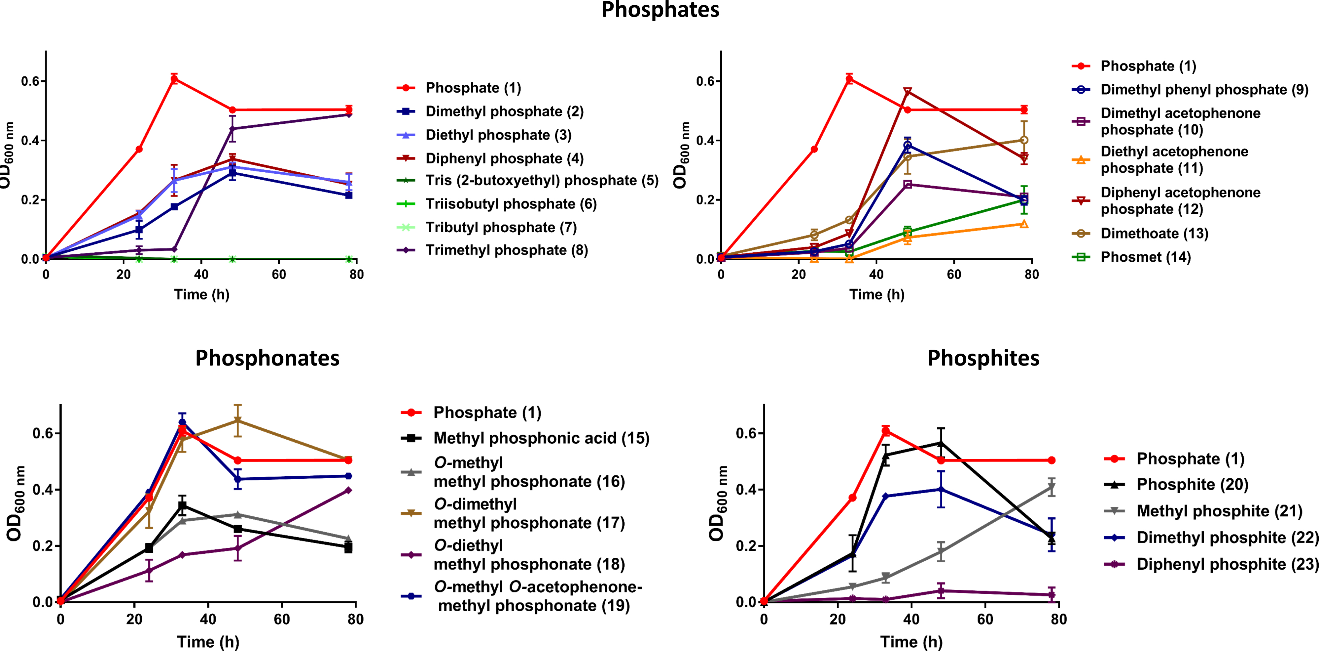

**Fig. S1: Growth curves of *R. pomeroyi* DSS3.** Artificial seawater medium was supplemented with 1 mM various organophosphate compounds (phosphodiesters, phosphotriesters, phosphonate mono- and diesters, phosphite mono- and diesters) as the sole phosphorous source. The growth was monitored over a period of 78 h. The highest OD values are presented in the heat map of **Fig. 3**.

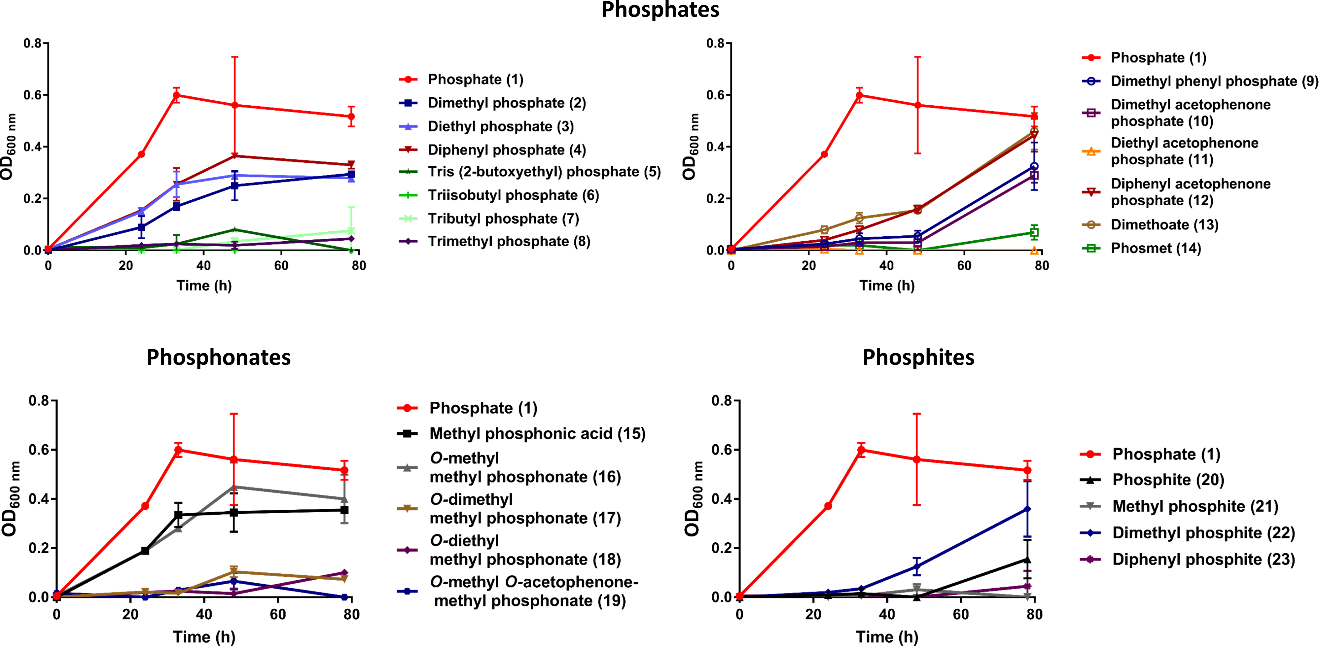

**Fig. S2: Growth curves of *Ruegeria sp.* TM1040.** Artificial seawater medium was supplemented with 1 mM various organophosphate compounds (phosphodiesters, phosphotriesters, phosphonate mono- and diesters, phosphite mono- and diesters) as the sole phosphorous source. The growth was monitored over a period of 78 h. The highest OD values are presented in the heat map of **Fig. 3**.

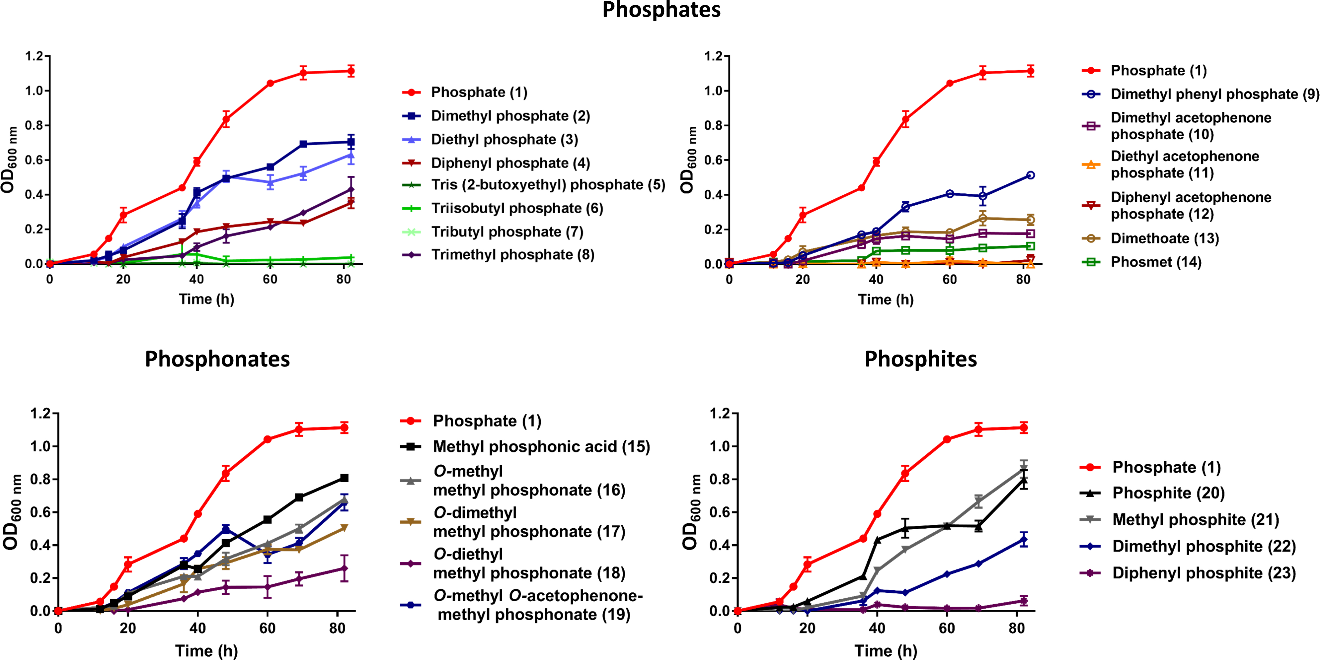

**Fig. S3: Growth curves of *P. inhibens*.** Artificial seawater medium was supplemented with 1 mM various organophosphate compounds (phosphodiesters, phosphotriesters, phosphonate mono- and diesters, phosphite mono- and diesters) as the sole phosphorous source. The growth was monitored over a period of 78 h. The highest OD values are presented in the heat map of **Fig. 3**.

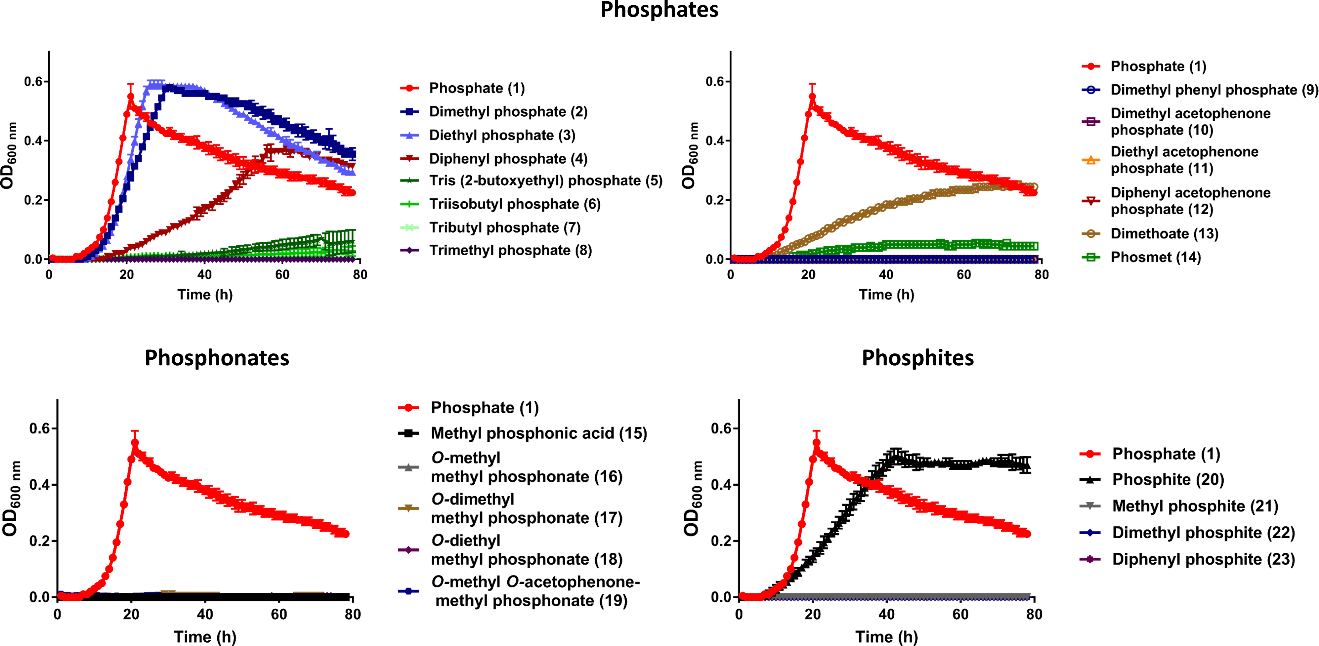

**Fig. S4: Growth curves of *Dinoroseobacter shibae.*** Artificial seawater medium was supplemented with 1 mM various organophosphate compounds (phosphodiesters, phosphotriesters, phosphonate mono- and diesters, phosphite mono- and diesters) as the sole phosphorous source. The growth was monitored over a period of 78 h. The highest OD values are presented in the heat map of **Fig. 3**.

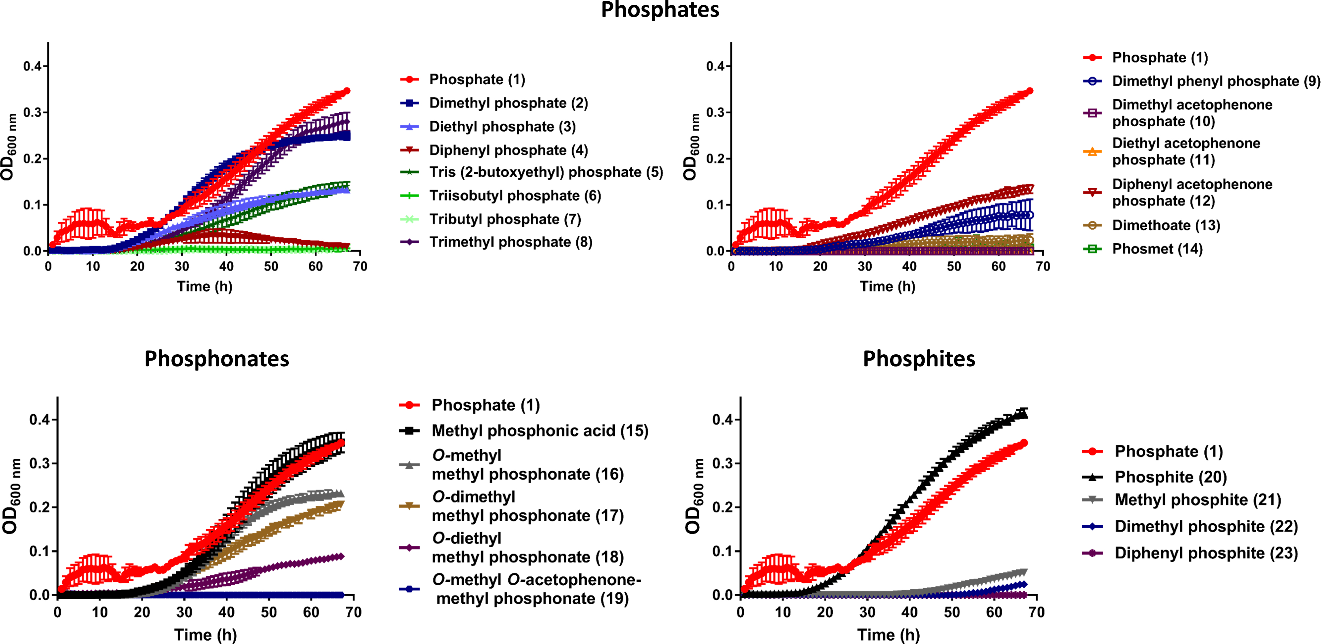

**Fig. S5: Growth curves of *Tateyamaria omphalii.*** Artificial seawater medium was supplemented with 1 mM various organophosphate compounds (phosphodiesters, phosphotriesters, phosphonate mono- and diesters, phosphite mono- and diesters) as the sole phosphorous source. The growth was monitored over a period of 67 h. The highest OD values are presented in the heat map of **Fig. 3**.

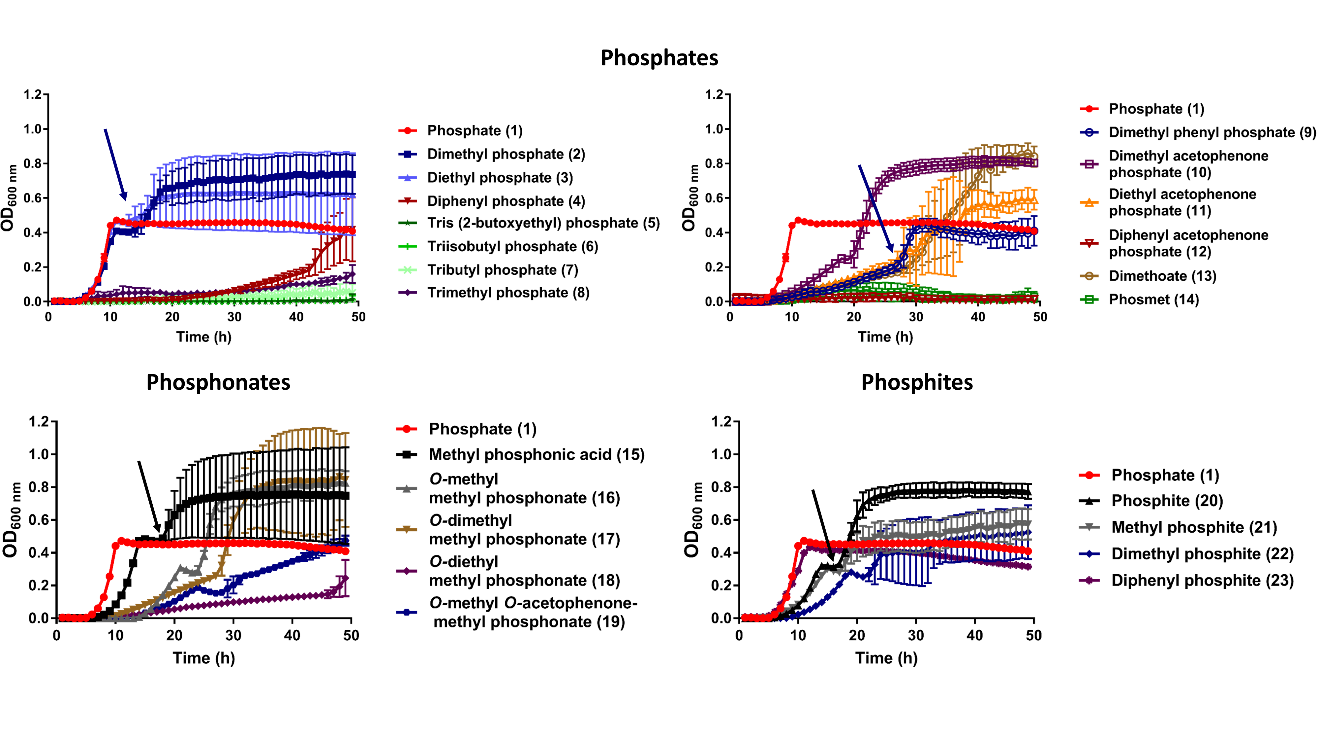

**Fig. S6: Growth curves of *Celeribacter naphtalenivorans.*** Artificial seawater medium was supplemented with 1 mM various organophosphate compounds (phosphodiesters, phosphotriesters, phosphonate mono- and diesters, phosphite mono- and diesters) as the sole phosphorous source. The growth was monitored over a period of 49 h. The arrows indicate the second growth peak which is possibly due to culture precipitation in the microtiter plate. Therefore, values for generating the heat map in **Fig. 3** were taken before the appearance of the second peak.

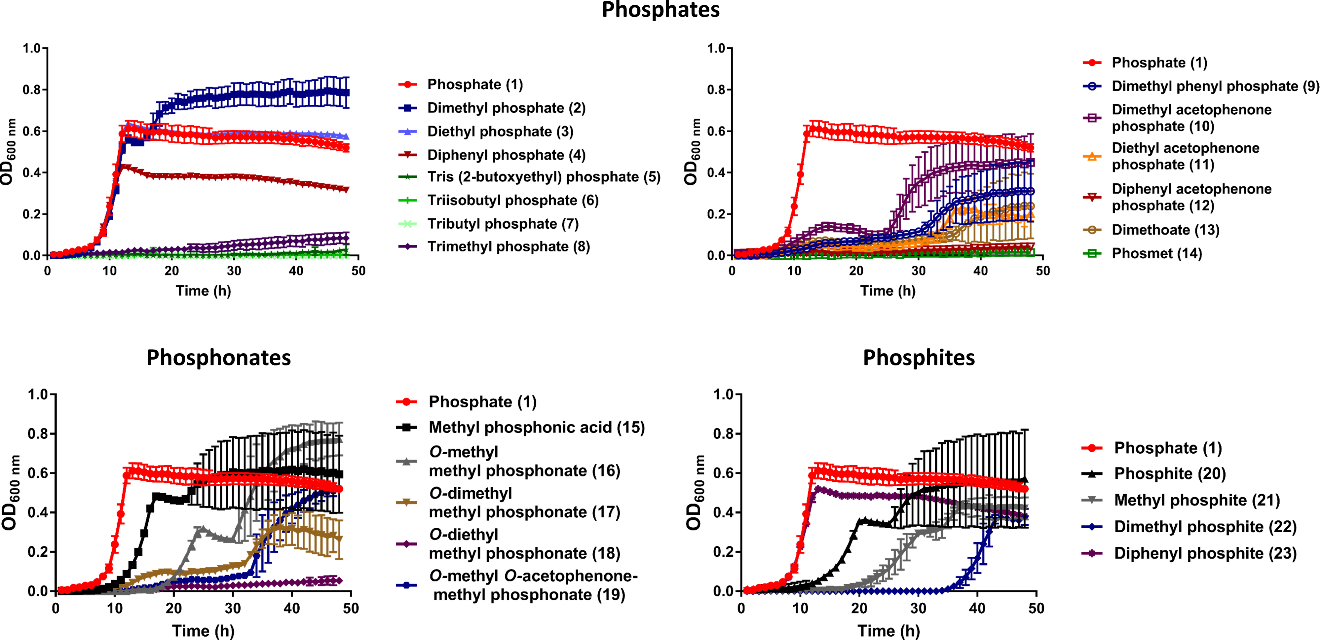

**Fig. S7: Growth curves of *Alteromononas macleodii*.** Artificial seawater medium was supplemented with 1 mM various organophosphate compounds (phosphodiesters, phosphotriesters, phosphonate mono- and diesters, phosphite mono- and diesters) as the sole phosphorous source. The growth was monitored over a period of 49 h. The max OD values for generating the heat map in **Fig. 3** were taken before the appearance of the second peak.

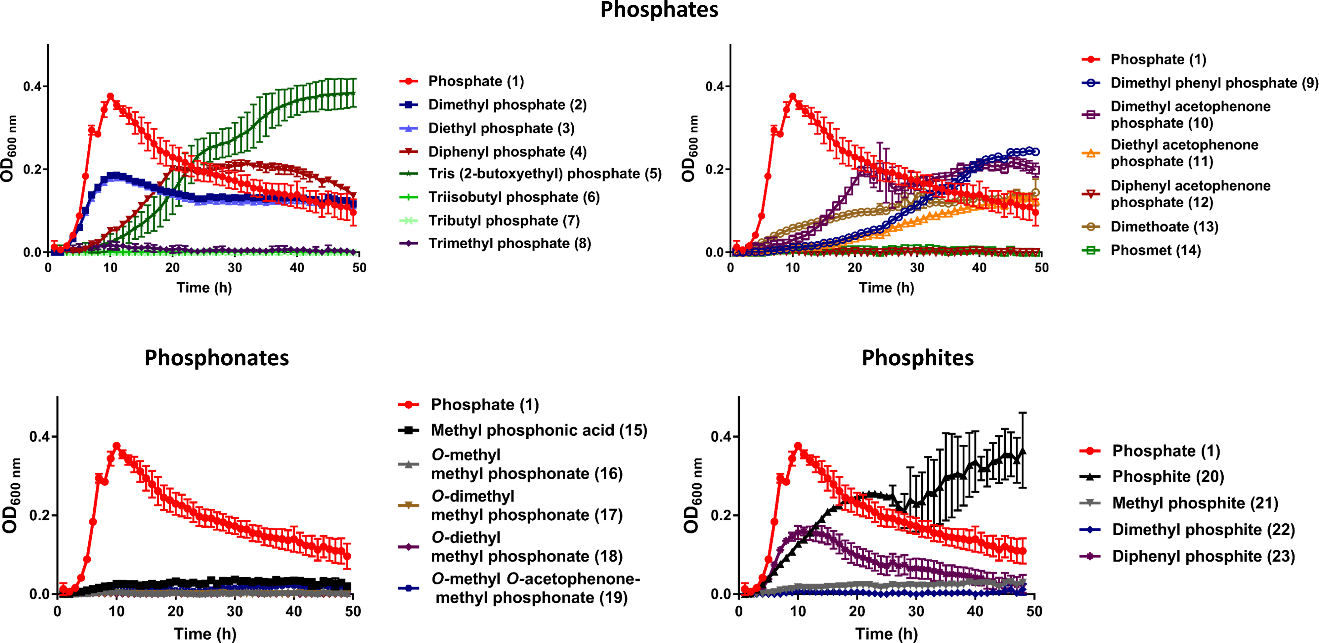

**Fig. S8: Growth curves of *Alteromononas mediterranea*.** Artificial seawater medium was supplemented with 1 mM various organophosphate compounds (phosphodiesters, phosphotriesters, phosphonate mono- and diesters, phosphite mono- and diesters) as the sole phosphorous source. The growth was monitored over a period of 49 h. The max OD values for generating the heat map in **Fig. 3** were taken before the appearance of the second peak.

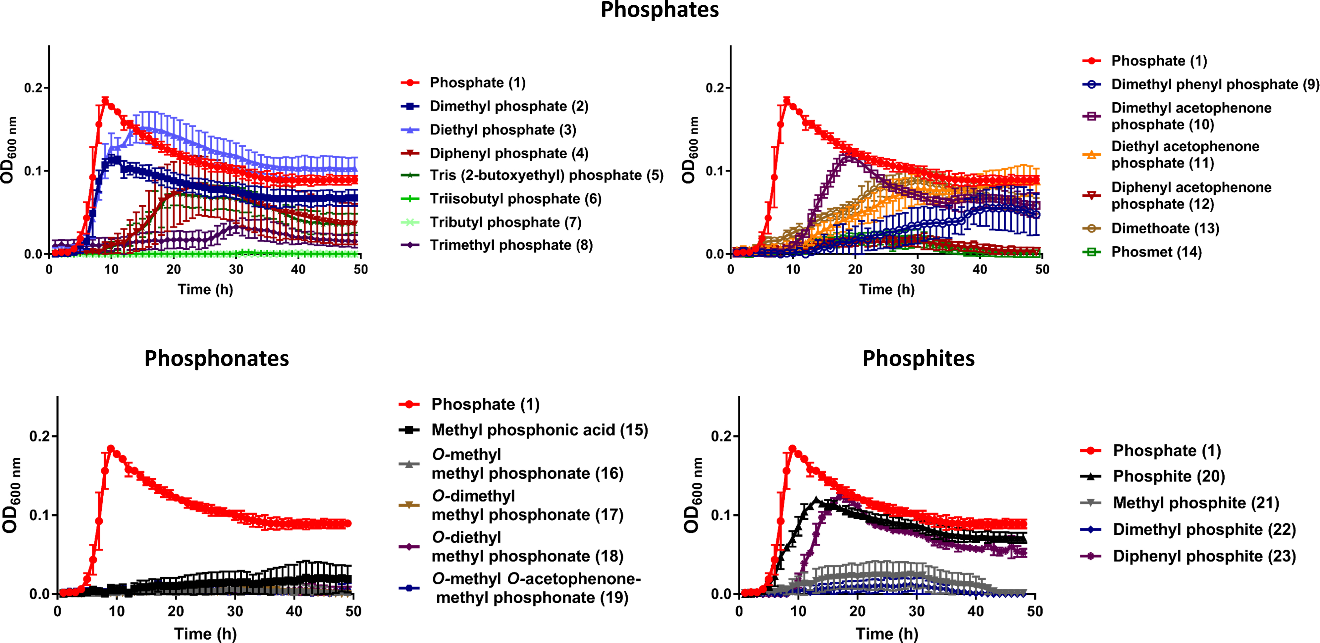

**Fig. S9: Growth curves of *Pseudoalteromonas sp.*.** Artificial seawater medium was supplemented with 1 mM various organophosphate compounds (phosphodiesters, phosphotriesters, phosphonate mono- and diesters, phosphite mono- and diesters) as the sole phosphorous source. The growth was monitored over a period of 49 h. the arrow. The max OD values for generating the heat map in **Fig. 3** were taken before the appearance of the second peak.

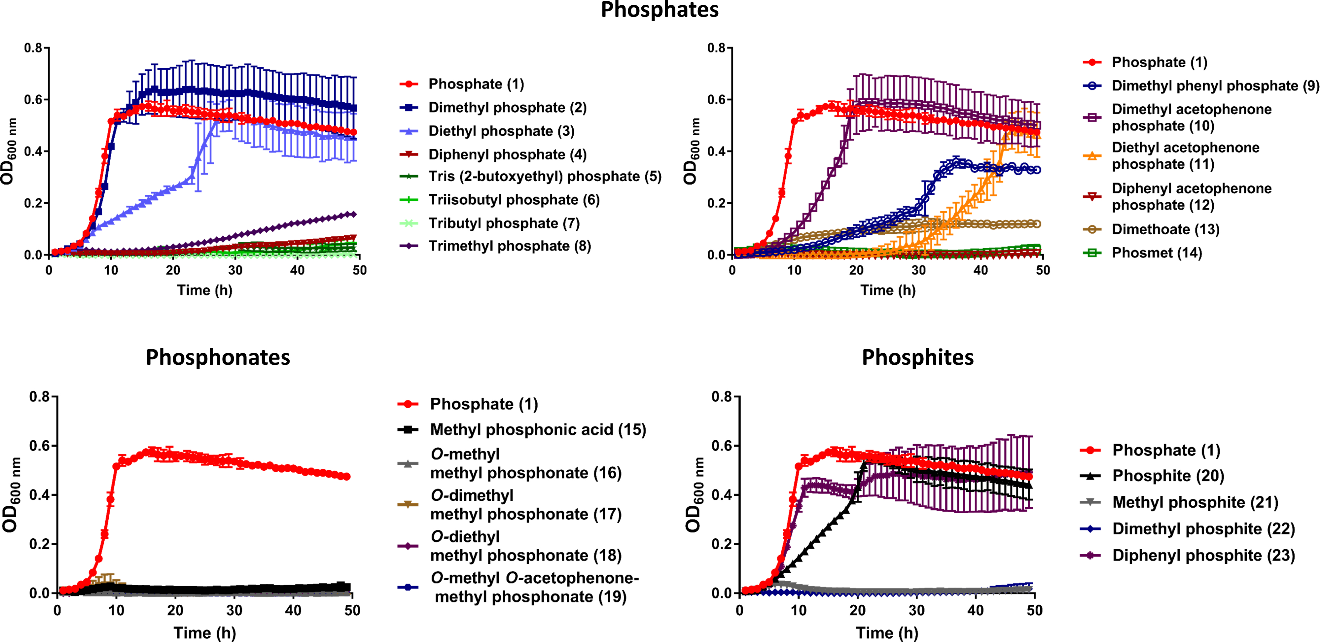

**Fig. S10: Growth curves of *Marinobacter sp.*.** Artificial seawater medium was supplemented with 1 mM various organophosphate compounds (phosphodiesters, phosphotriesters, phosphonate mono- and diesters, phosphite mono- and diesters) as the sole phosphorous source. The growth was monitored over a period of 49 h. The max OD values for generating the heat map in **Fig. 3** were taken before the appearance of the second peak.

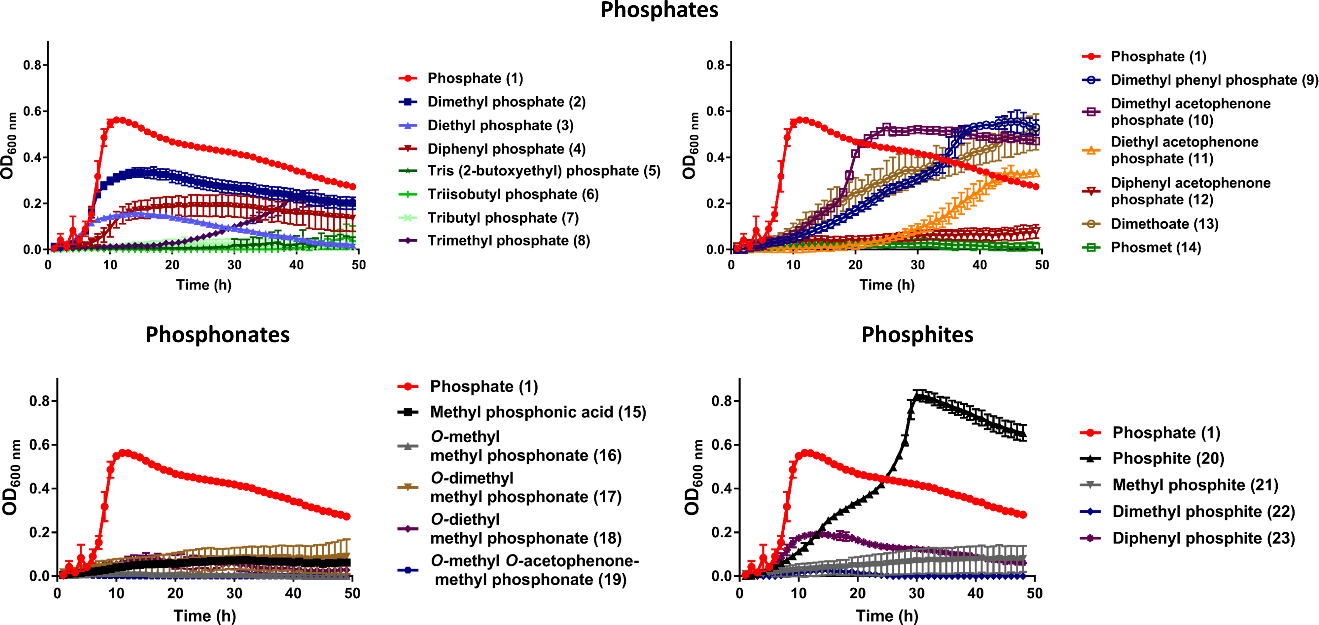

**Fig. S11: Growth curves of *Cobetia sp.*.** Artificial seawater medium was supplemented with 1 mM various organophosphate compounds (phosphodiesters, phosphotriesters, phosphonate mono- and diesters, phosphite mono- and diesters) as the sole phosphorous source. The growth was monitored over a period of 49 h. The max OD values for generating the heat map in **Fig. 3** were taken before the appearance of the second peak.

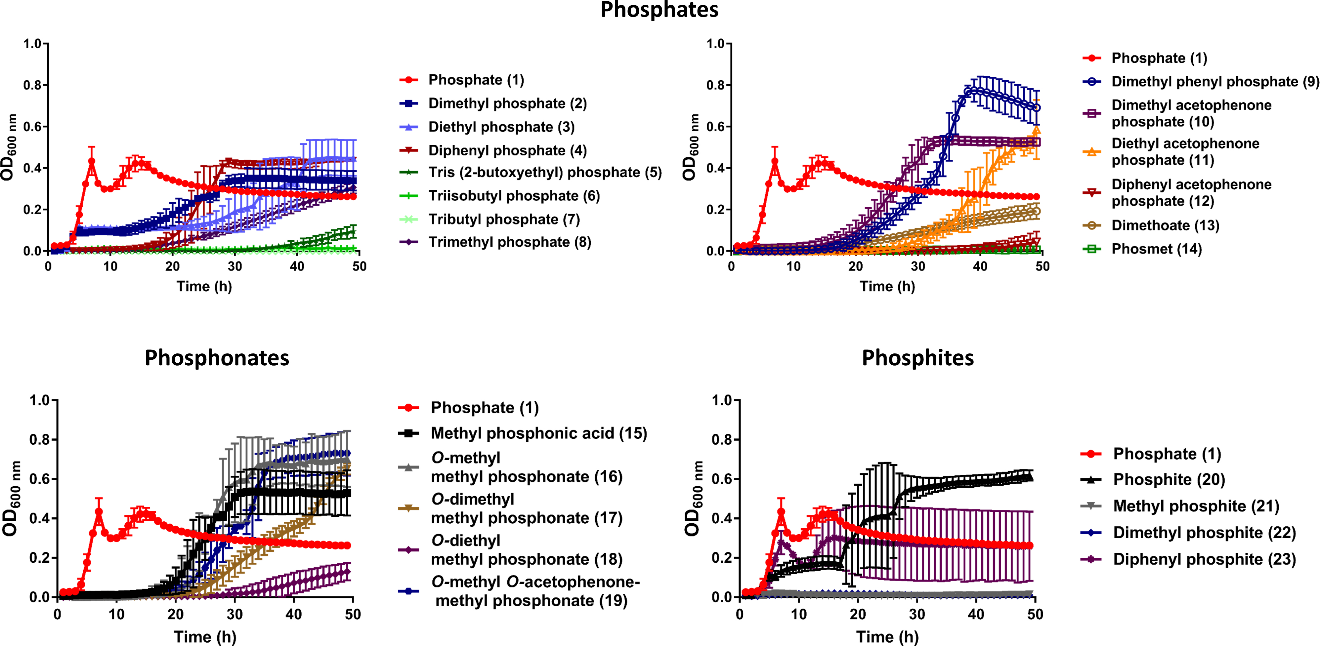

**Fig. S12: Growth curves of *Marinomonas brasilensis*.** Artificial seawater medium was supplemented with 1 mM various organophosphate compounds (phosphodiesters, phosphotriesters, phosphonate mono- and diesters, phosphite mono- and diesters) as the sole phosphorous source. The growth was monitored over a period of 49 h. The max OD values for generating the heat map in **Fig. 3** were taken before the appearance of the second peak.

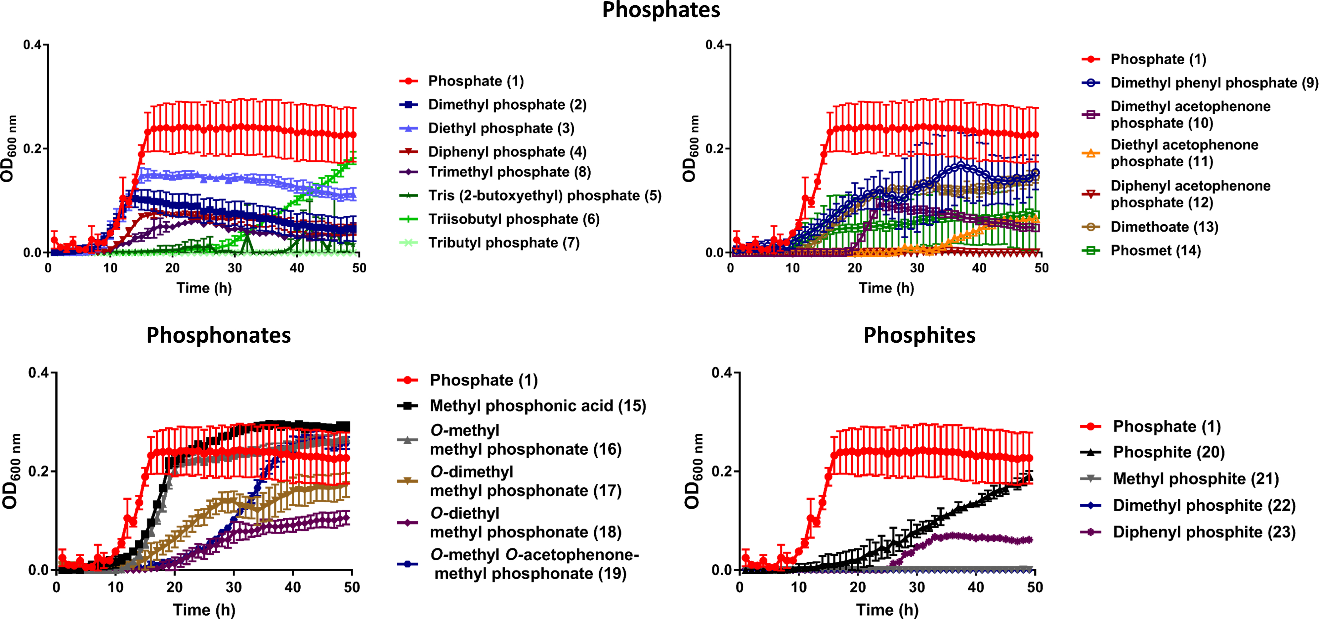

**Fig. S13: Growth of *Labrenzia sp.*.** Artificial seawater medium was supplemented with 1 mM various organophosphate compounds (phosphodiesters, phosphotriesters, phosphonate mono- and diesters, phosphite mono- and diesters) as the sole phosphorous source. The growth was monitored over a period of 49 h. the arrow. The max OD values for generating the heat map in **Fig. 3** were taken before the appearance of the second peak.

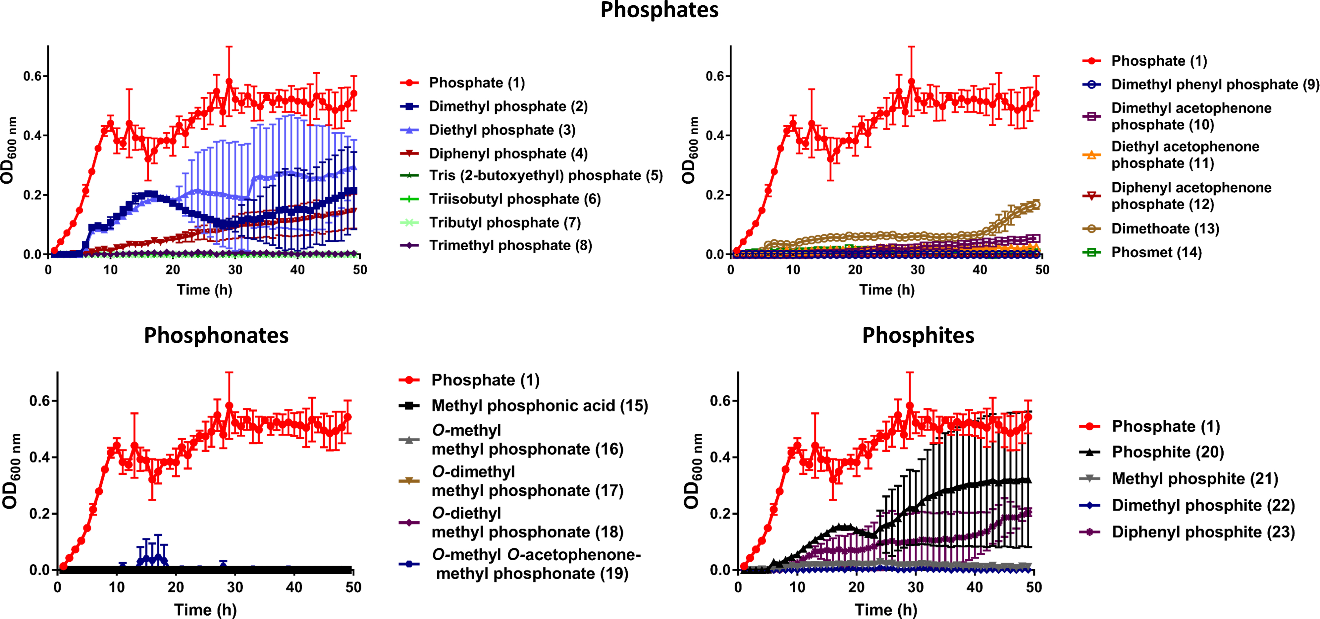

**Fig. S14: Growth of the *Vibrio sp.*.** Artificial seawater medium was supplemented with 1 mM various organophosphate compounds (phosphodiesters, phosphotriesters, phosphonate mono- and diesters, phosphite mono- and diesters) as the sole phosphorous source. The growth was monitored over a period of 49 h. The max OD values for generating the heat map in **Fig. 3** were taken before the appearance of the second peak.

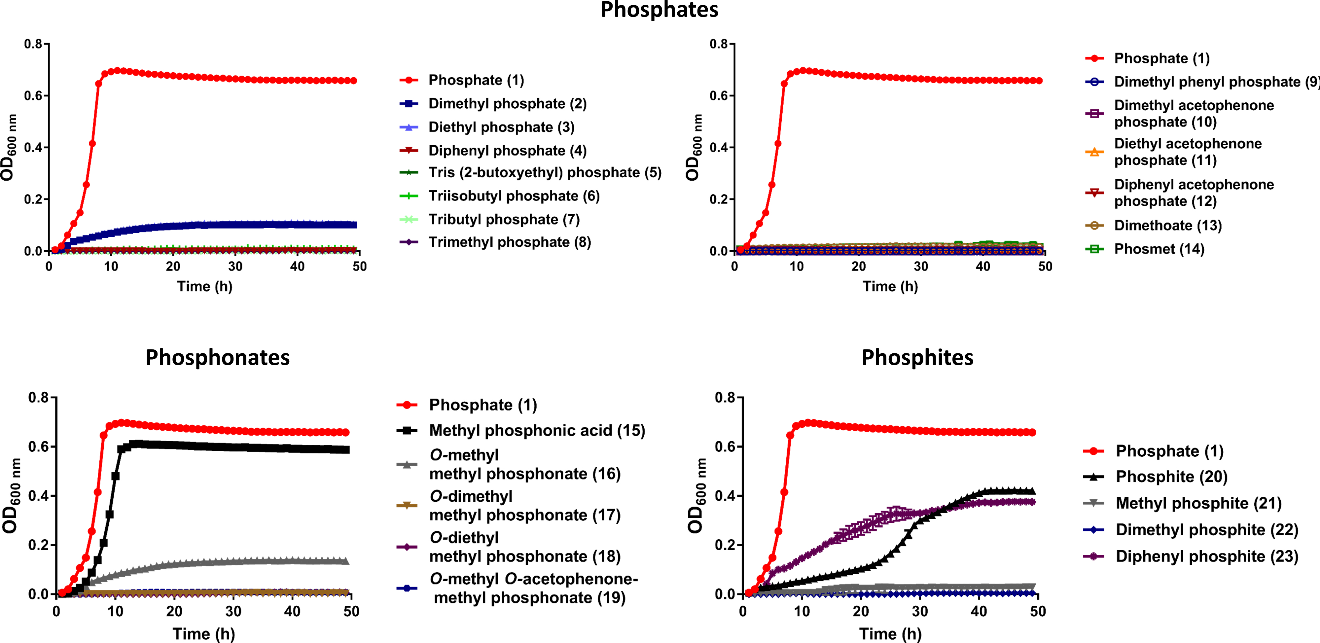

**Fig. S15: Growth curves of *E. coli* BL21.** MOPS medium was supplemented with 1 mM various organophosphate compounds (phosphodiesters, phosphotriesters, phosphonate mono- and diesters, phosphite mono- and diesters) as the sole phosphorous source. The growth was monitored over a period of 49 h. The highest OD values are presented in the heat map of **Fig. 3**.

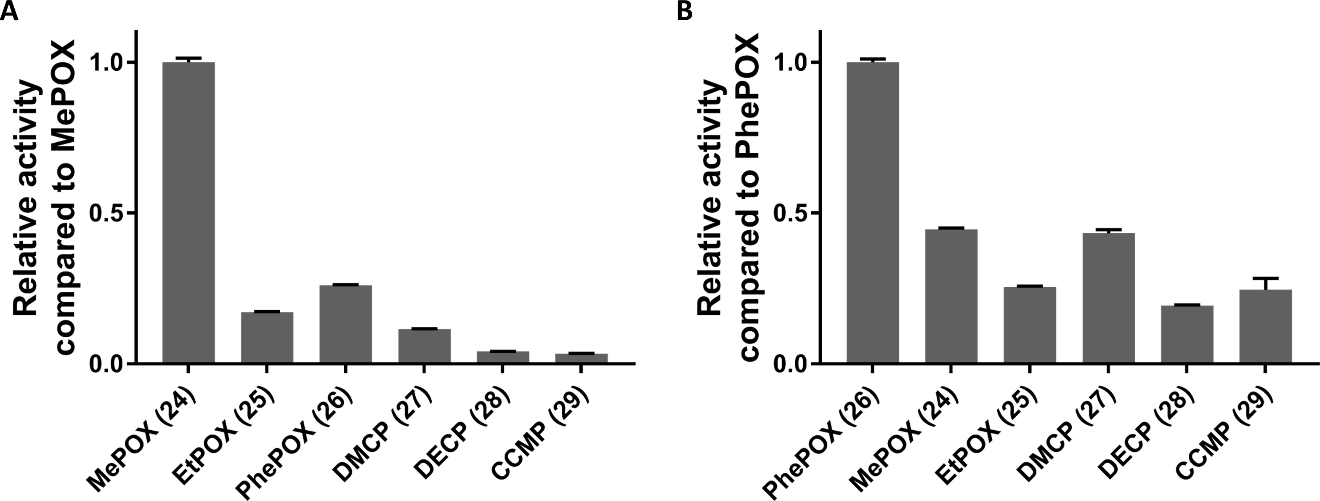

**Fig. S16: Activity profiling of *R. pomeroyi* DSS3 (A) and *Ruegeria sp.* TM1040 (B).** Lysates of both strains were incubated in 100 mM Tris pH 8.0, 50 mM NaCl with 1 mM substrates (see **Table S2**). The increase of absorbance at 405 nm was measured and values were normalized according to the best utilized substrate whose activity was set to 1.

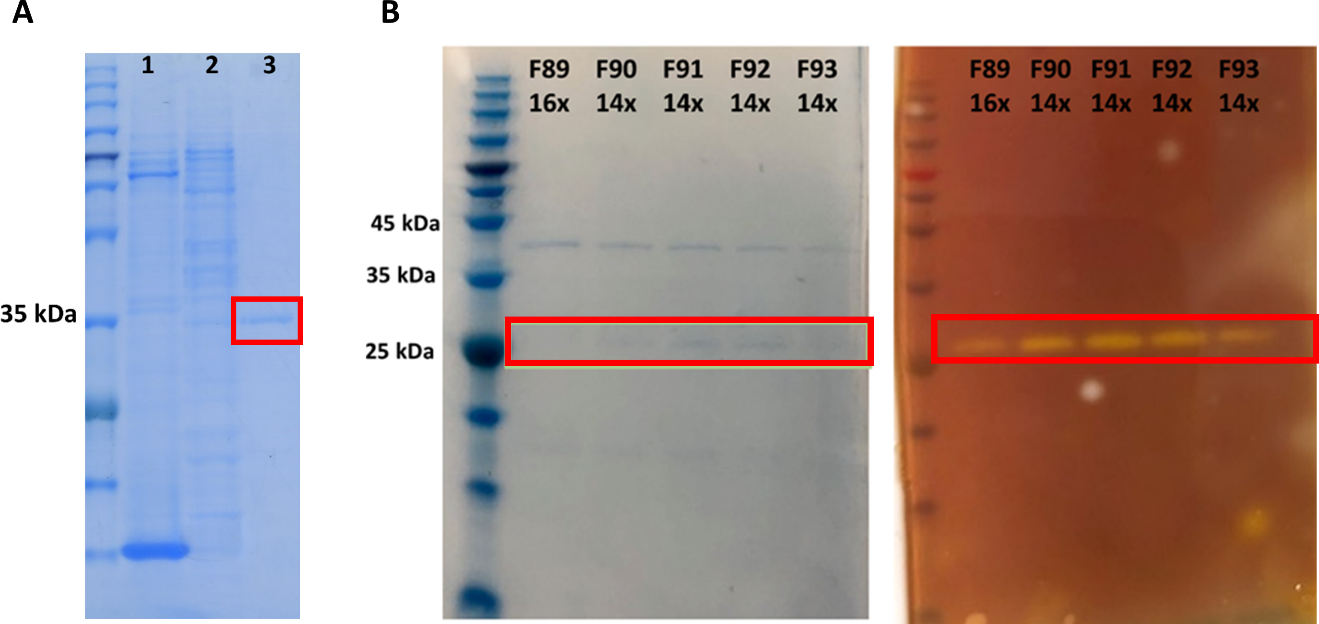

**Fig. S17: Isolation and identification of phosphotriesterases from *Ruegeria sp.* TM1040 and *R. pomeroyi* DSS3. A** SDS-PAGE analysis of *Ruegeria sp.* TM1040’s phosphotriesterase purification steps. 1- crude cell lysate, 2 – the most active fractions from anion exchange chromatography, 3 – the eluted fraction from the affinity matrix **B** *In situ* phosphotriesterase activity of the fractions after hydrophobic butyl chromatography. Active fractions were concentrated 14-16-fold and 10 µl was loaded on SDS Page. The first half of the gel was stained with Commassie blue (**left)** and the second half was placed on 1.5% agar gel with 100 mM NaCl pH 8.0, 0.4 mg/ml Cresol purple, 1 mM methyl-paraoxon (**right**). After few minutes yellow color was developed slightly above 25 kDa protein marker. Samples labeled in red were cut out from the gel **(A)** and send for enzyme identification by mass spectrometry or pulled together and then re-analyzed by SDS-PAGE and active band sent for identification by mass spectrometry **(B)**.

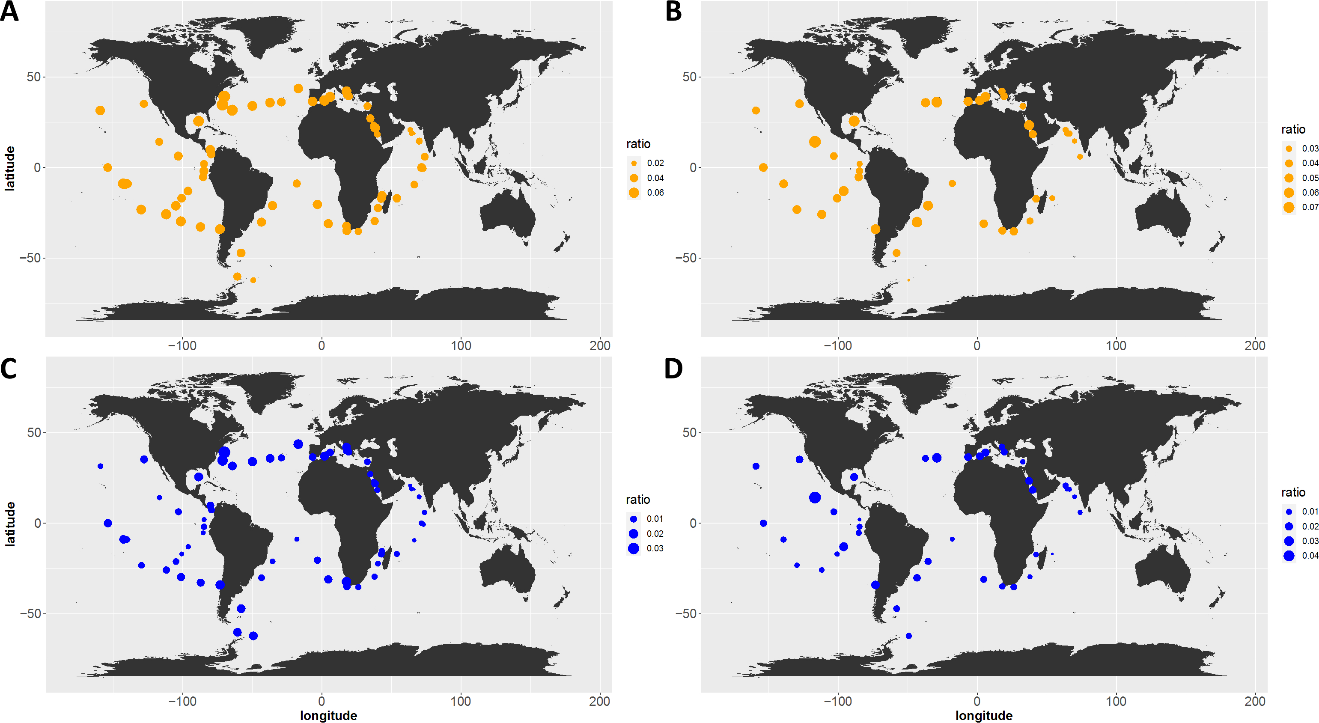

**Fig. S18: Matallo beta-lactamase PTE (A, B) and cyclase-PTE (C, D) genes abundances in the TARA Oceans dataset.** For both cyclase-PTE and MBL-PTE genes the abundances (percent of total reads) were identified in the TARA Oceans prokaryotic dataset using Ocean Gene Atlas in the surface waters (SRF, panels **A** and **C**) and deep chlorophyll maximum (DCM, panels **B** and **D**) for all fraction sizes. The values are presented as the ratio between the identified MBL-PTEs or cyclase-PTEs and the *recA* abundance per TARA Oceans sample.

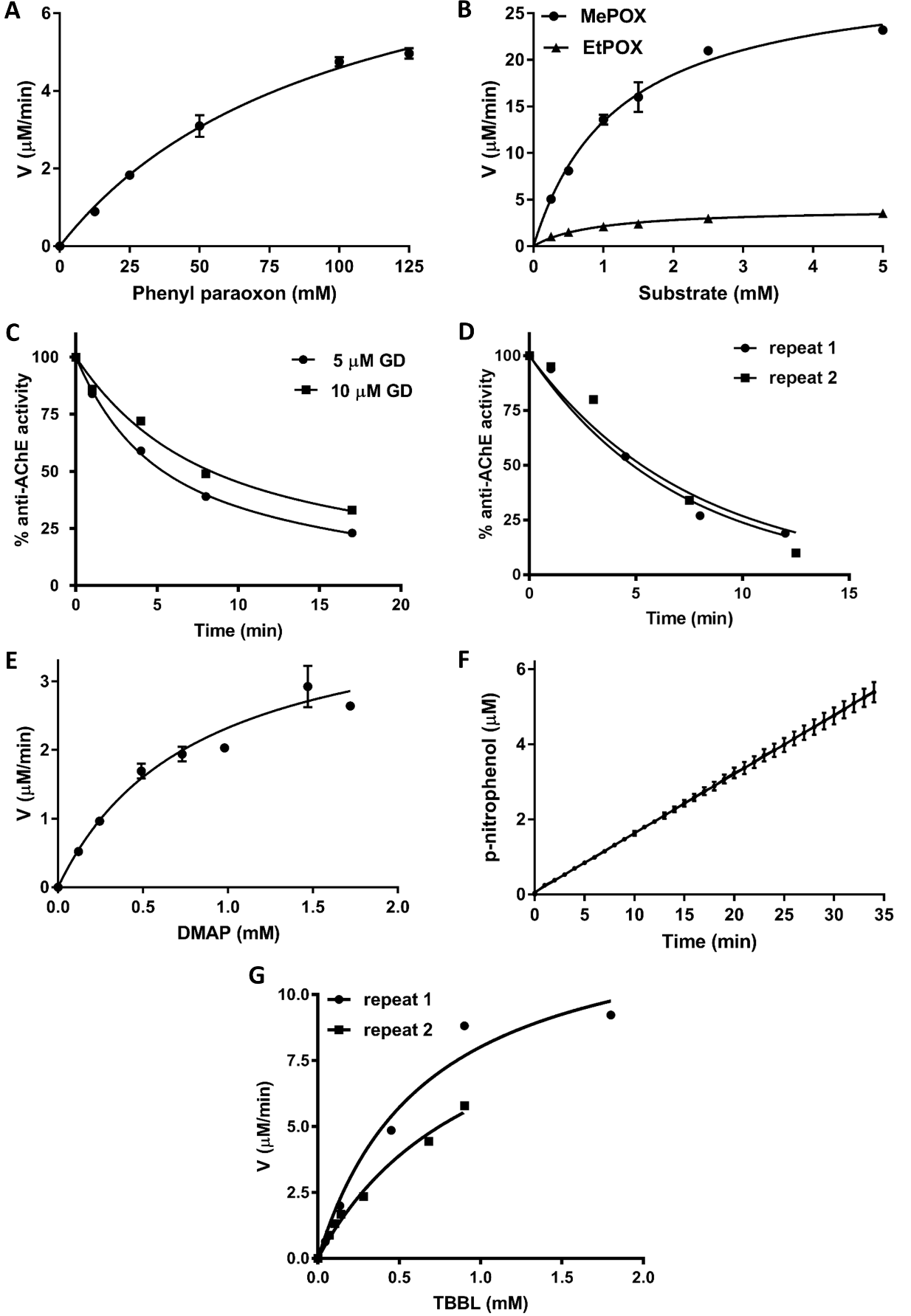

**Fig. S19: Kinetic characterization metallo beta-lactamase (MBL-PTE).** Data fit to Michaelis-Menten model for **A** phenyl paraoxon (PhPOX, compound 26) **B** methyl paraoxon (MePOX, compound 24) and paraoxon (EtPOX, compound 25); **E** dimethyl acetophenone phosphate (DMAP, compound 10) and **G** TBBL lacton (compound 35). **C** GD (compound 32) and **D** GF (compound 33) detoxification was monitored by prevention of acetylcholine esterase activity. **E** Hydrolysis of 0.95 mM methyl parathion (compound 31) in the presence of 0.84 µM MBL-PTE.

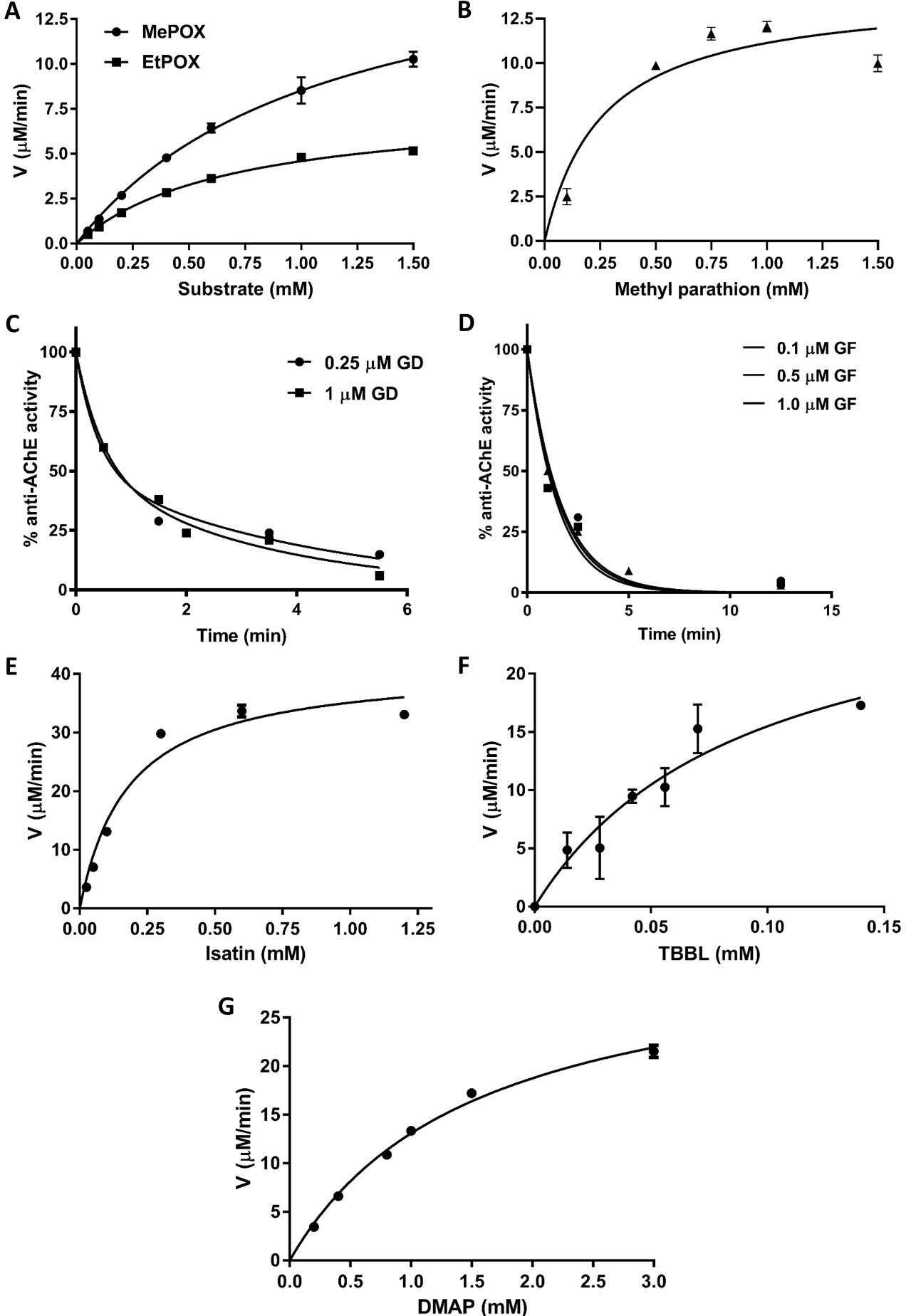
**Fig. S20: Kinetic characterization of cyclase-PTE.** Data fit to Michaelis-Menten model for **A** methyl paraoxon (MePOX, compound 24) and paraoxon (EtPOX, compound 25); **B** methyl parathion (compound 31); **E** isatin (compound 34); **F** TBBL lacton (compound 35) and **G** dimethyl acetophenone phosphate (DMAP, compound 10). **C** GD (compound 32) and **D** GF (compound 33) detoxification was monitored by prevention of acetylcholine esterase activity.

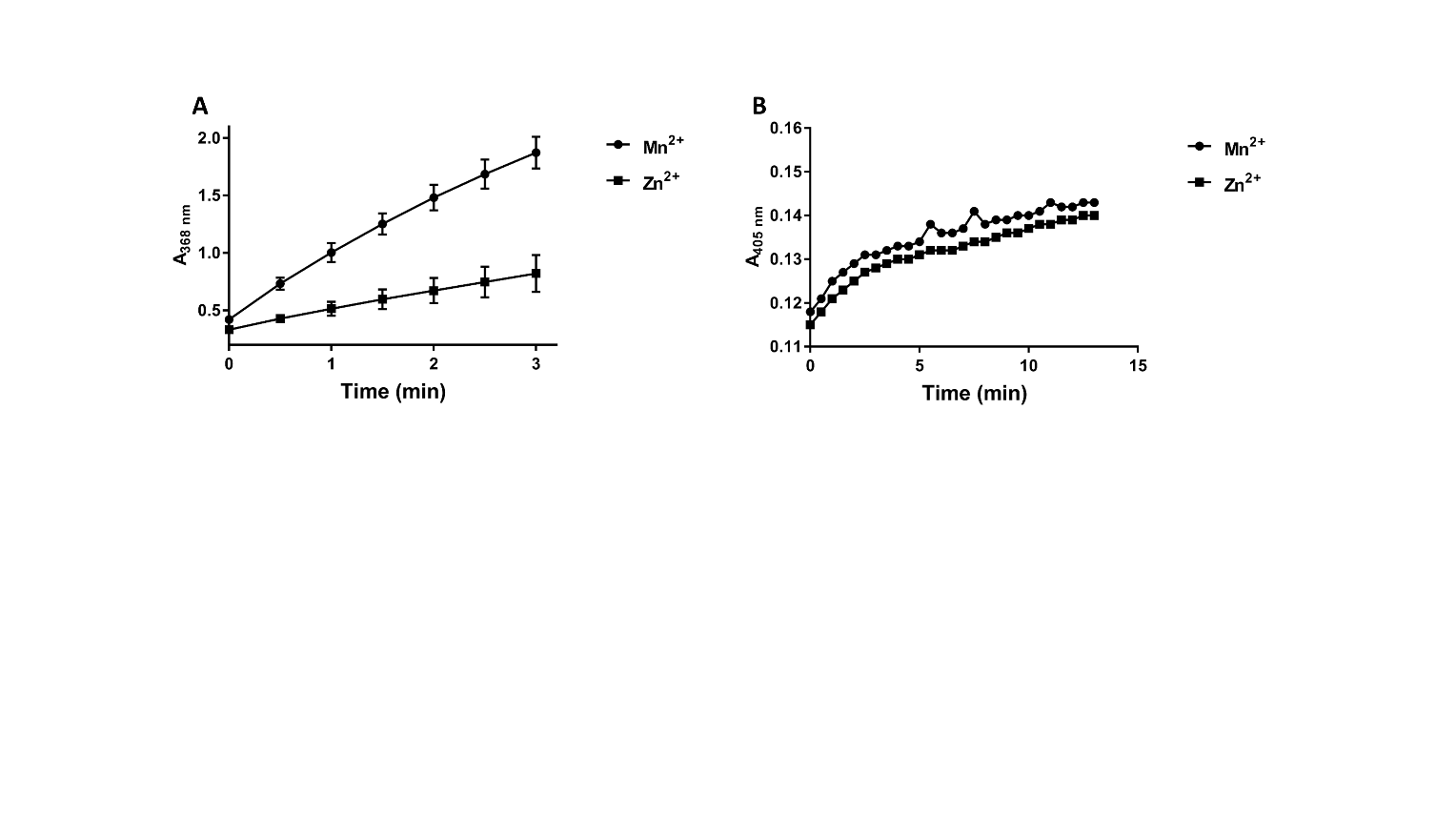

**Fig. S21: Phosphotriesterase and isatin hydrolase activity in the cell lysate of *E. coli* BL21 expressing isatin hydrolase A from *Labrenzia aggregate.* A** Isatin (compound 34) hydrolase activity in the cell lysate diluted 2000-fold in the reaction. **B** Methyl-paraoxon (compound 24) hydrolysis in the cell lysate diluted 10-fold in the reaction.

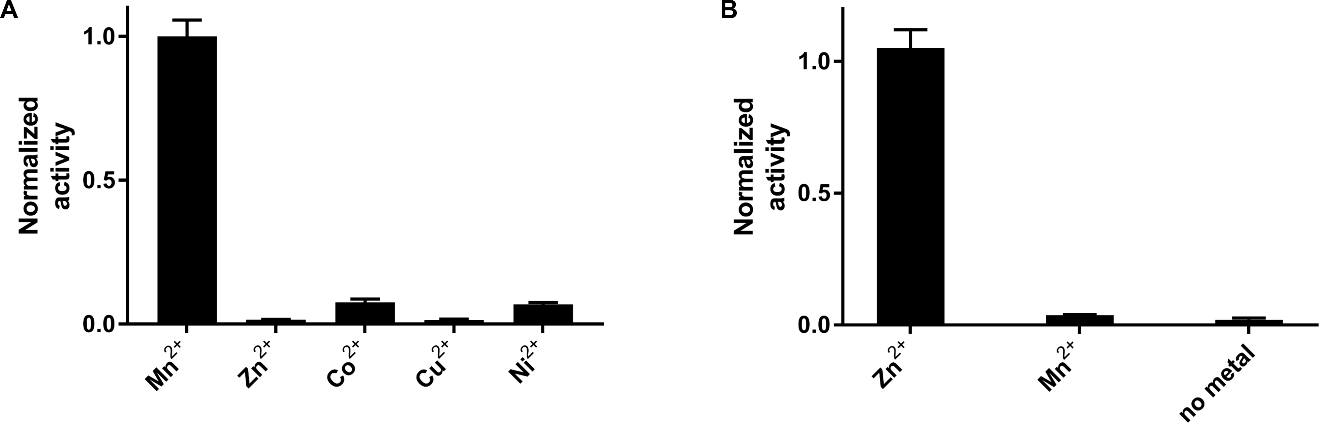

**Fig. S22: Identification of the catalytic metal of MBL-PTE (A) and cyclase-PTE (B).** Enzymes are expressed in the presence of various metals, and activity measured in the cell lysate with 0.1 mM phenyl paraoxon (compound 26, substrate for MBL-PTE) and 1 mM methyl paraoxon (compound 24, cyclase-PTE substrate). The release of p-nitrophenol at 405 nm was monitored, values were normalized according to the highest activity whose value was set to 1.

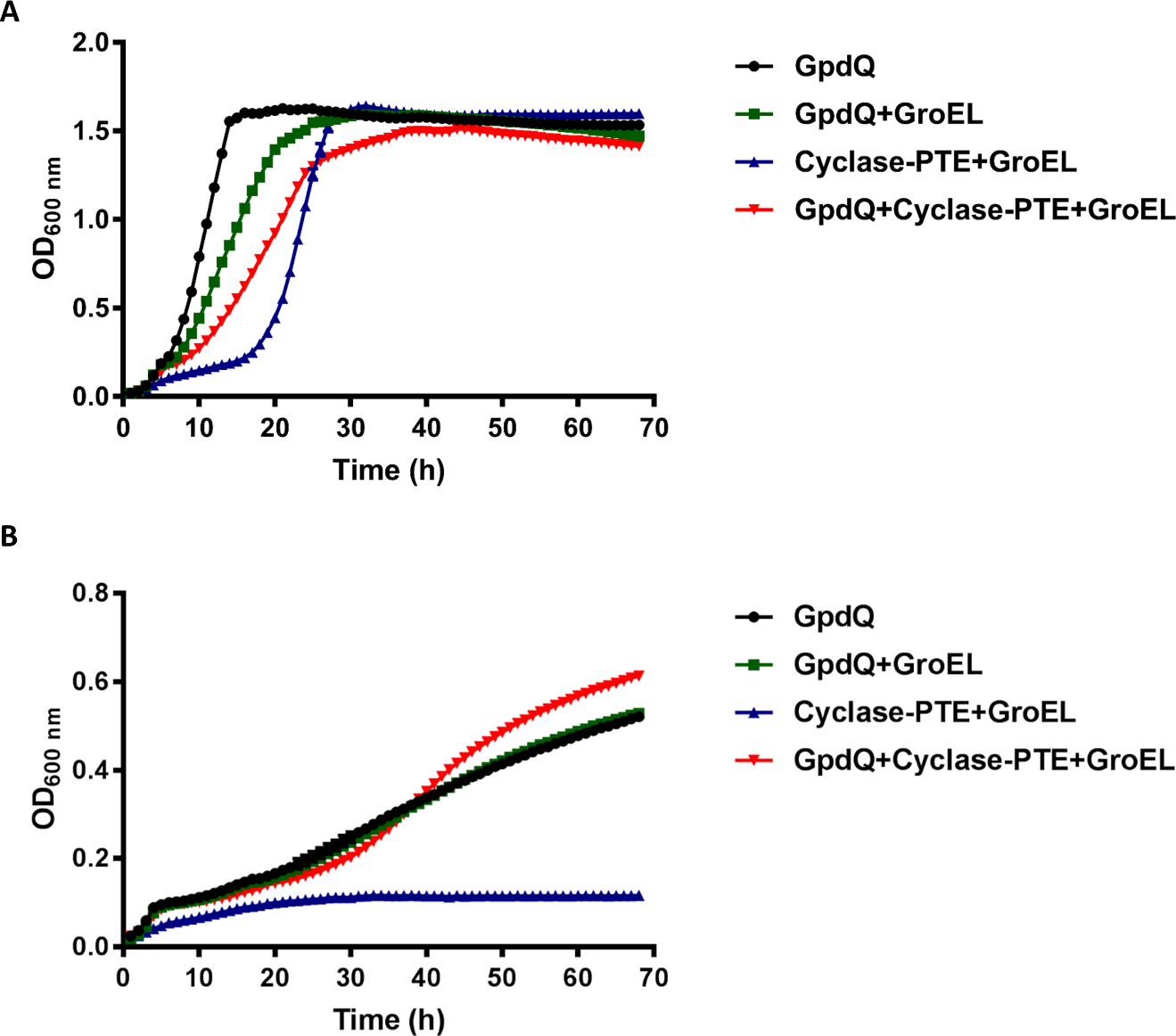

**Fig. S23: Growth of *E. coli* BL21 co-expressing GpdQ and cyclase-PTE on: (A) phosphate and (B) dimethyl acetophenone phosphate (DMAP, compound 10) as the sole phosphorous source.** The growth was monitored in MOPS media supplemented with 1 mM dimethyl acetophenone phosphate for 68 h at 30°C. Co-expression of cyclase-PTE and chaperones (GroEL) decreases the growth of *E. coli* BL21 in the phosphate rich medium **(A)**, the effect of cyclase-PTE on the growth of *E. coli* BL21 on dimethyl acetophenone phosphate **(B)** was significantly lower than expected possibly due to toxicity of the 3 genes co-expression.

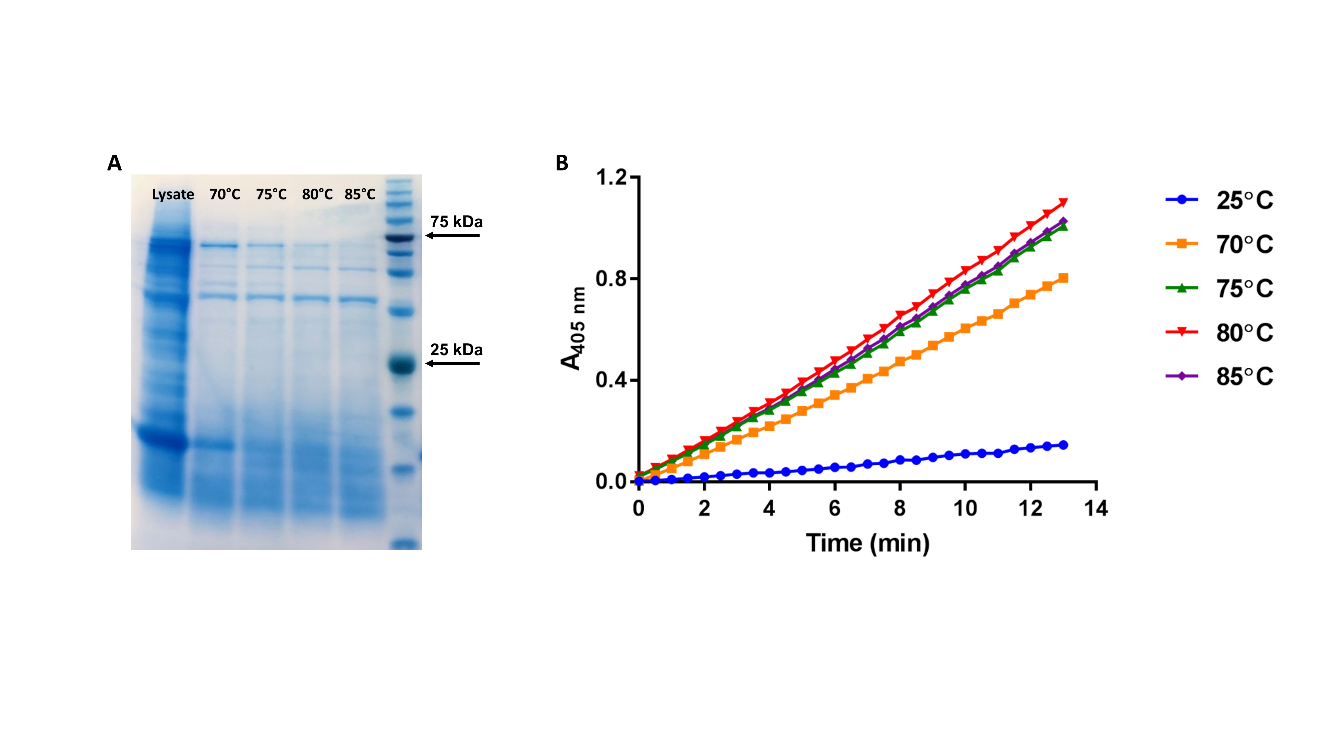

**Fig. S24: Heat activation of the PTE activity in *R. pomeroyi* DSS3 crude cell lysate.** The cell pellet was lysed in 50 mM Tris, 50 mM NaCl pH 8.0, 0.1% Triton X-100, 0.1% Tergitol NP-10, 0.2 mg/ml Lysozyme, 5 µl benzonase (≥250 units/μL, ≥90%), incubated for 30 min at various temperatures. **A** The lysate was analyzed by SDS-PAGE and **B** residual activity was measured with methyl-paraoxon (compound 24).

**Tables**

**Table S1: The list of organophosphate (OP) substrates used for growth profiling of various marine bacteria.** A number is associated with each compound, numbers are denoted throughout the text for convenience. Substrates used for profiling are divided into several groups: 1. Phosphotriesters and products of their hydrolysis pathway phosphodiesters and phosphomonoesters. Phosphotriesters are divided into 4 categories: commonly used flame retardants (5, 6, 7), phosphotriesters that differ in: the acidity of the leaving group (8, 9, 10) and size of the substituent groups (10, 11, 12) and the last group are insecticides (13, 14). 2. Phosphonate diesters (17, 18, 19) and the products of their hydrolysis pathway (15 and 16). Phosphite diesters (22 and 23) and the products of their hydrolysis pathway (20 and 21).

| **Phosphotriesters** | | | | | | | |
| --- | --- | --- | --- | --- | --- | --- | --- |
| **Commonly used** | | | **Leaving group p*Ka*** | | | **Size of substituents** | |
| 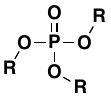  **R:**   - 2-butoxyethyl **(5)** - isobutyl **(6)** - butyl **(7)** | | | 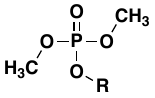  **R:**   - methyl **(8)** - phenyl **(9)** - acetophenone **(10)** | | | 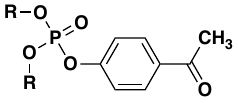  **R:**   - methyl **(10)** - ethyl **(11)** - phenyl **(12)** | |
| **Phosphodiesters** | | | **Insecticides (Thiophosphates)** | | | | |
| 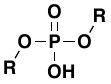  **R:**   - methyl **(2)** - ethyl **(3)** - phenyl **(4)** | | | 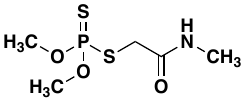  Dimethoate **(13)** | | 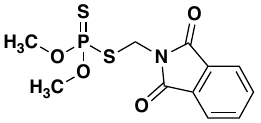  Phosmet **(14)** | | |
| **Phosphonate/phosphite monoesters** | | | | **Phosphonate/phosphite diesters** | | | |
| 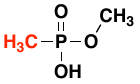  **(16)** | | 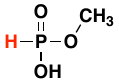  **(21)** | | 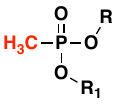  **R, R_1_:**   - methyl **(17)** - ethyl **(18)** - R=methyl,   R^1^=acetophenone **(19)** | | | 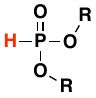  **R:**   - methyl **(22)** - phenyl **(23)** |
| **Other** | Phosphate **(1)**, Methyl phosphonic acid **(15)**, Phosphite **(20)** | | | | | | |
| **Structures** | 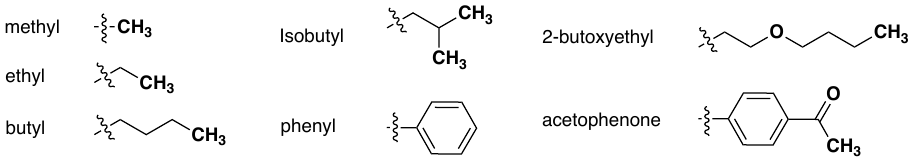 | | | | | | |

**Table S2: Substrates used for lysate profiling of *R. pomeroyi* DSS3 and *Ruegeria sp.* TM1040 and characterization of cyclase-PTE and MBL-PTE.** Diethyl p-nitrophenyl phosphate (compound 25) is generally labeled as paraoxon.

| **Name** | **Structure** |
| --- | --- |
| Dimethyl p-nitrophenyl phosphate  (Methyl paraoxon, MePOX)  (24) | 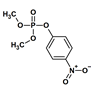 |
| Diethyl p-nitrophenyl phosphate  (Paraoxon, EtPOX)  (25) | 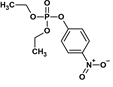 |
| Diphenyl p-nitrophenyl phosphate  (Phenyl paraoxon, PhePOX)  (26) | 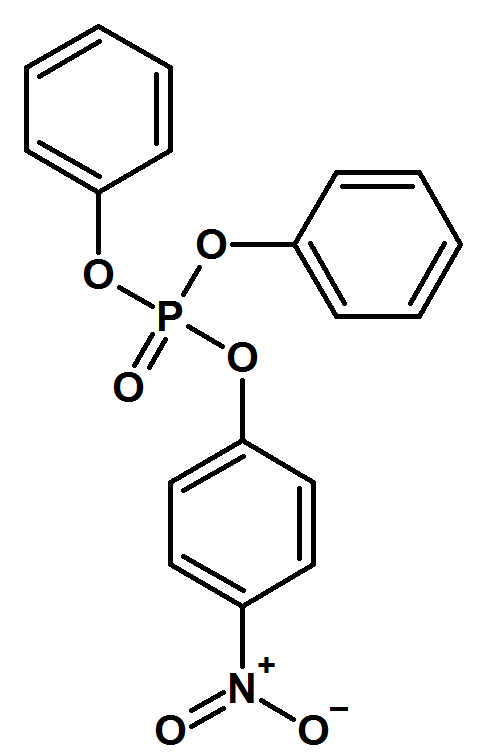 |
| Dimethyl coumarin phosphate  (DMCP)  (27) | 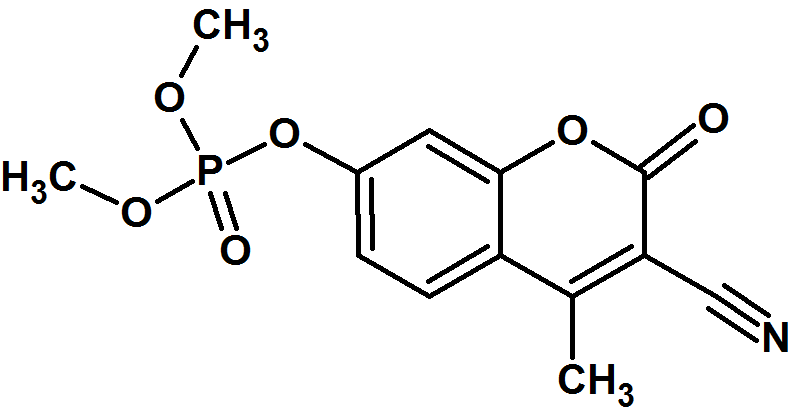 |
| Diethyl coumarin phosphate  (DECP)  (28) | 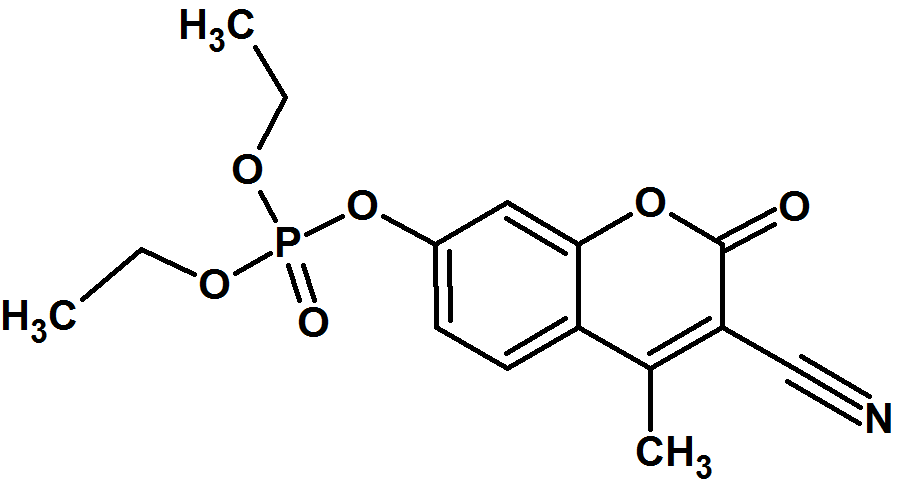 |
| *O-*cyclohexyl *O-*coumarin methyl phosphonate  (CCMP)  (29) | 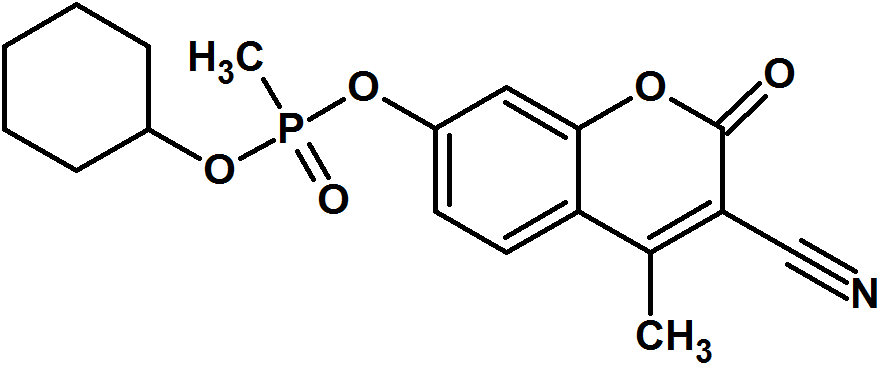 |
| Diethyl thio p-nitrophenyl phosphate  (Parathion, EtPTH)  (30) | 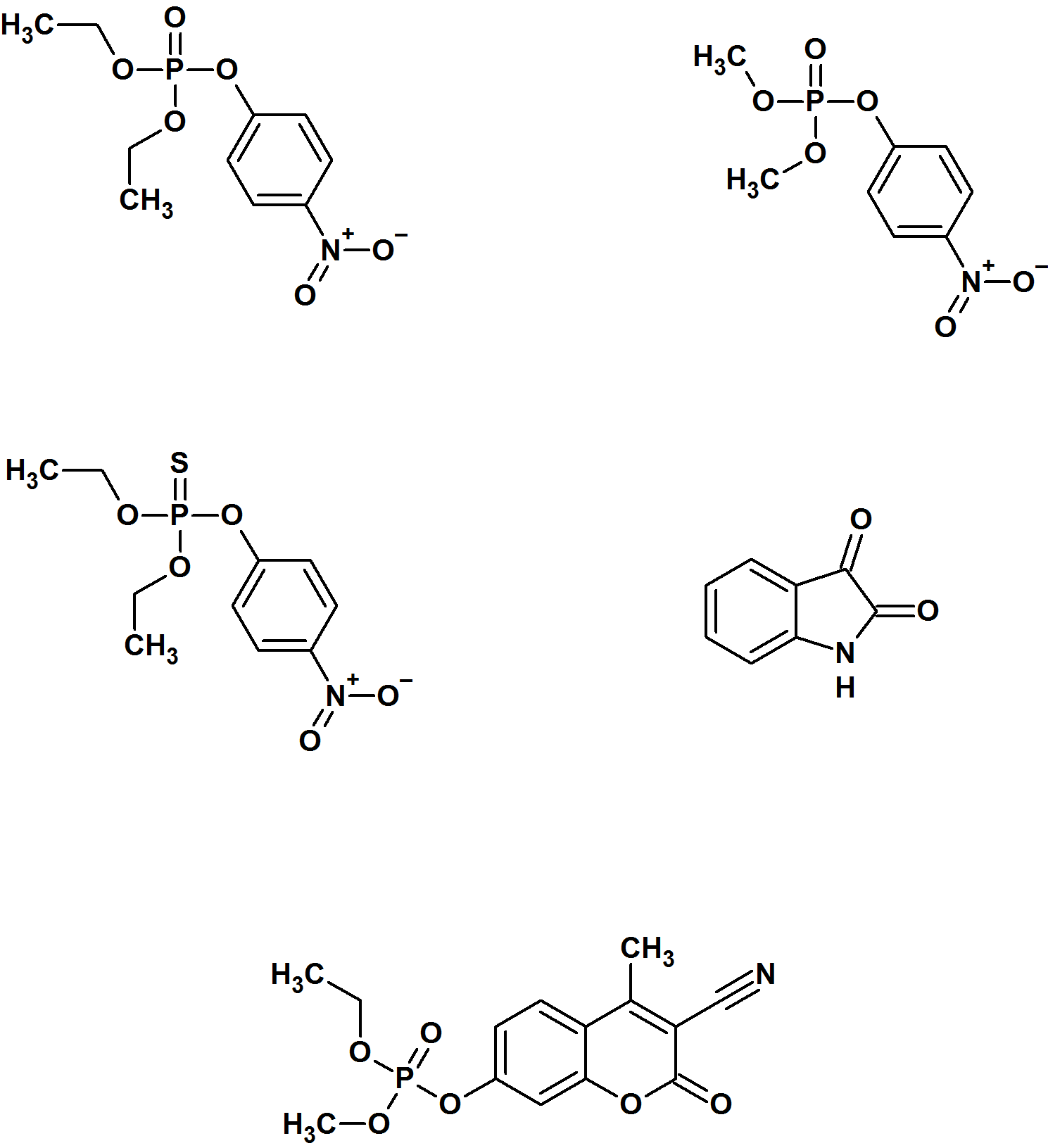 |
| Dimethyl thio p-nitrophenyl phosphate  (Methyl parathion, MePTH)  (31) | 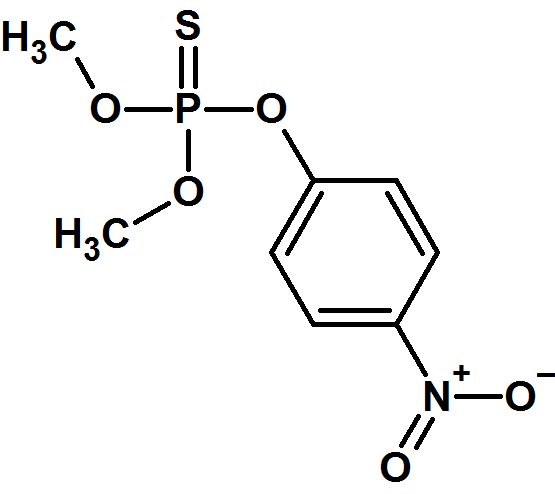 |
| (*SpR_C_/SpS_C_*)-soman  (GD)  (32) | 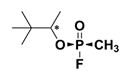 |
| (*Sp)*-cyclosarin  (GF)  (33) | 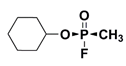 |
| Isatin  (34) | 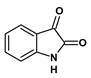 |
| Thiobutyl butyrolactone  (TBBL)  (35) | 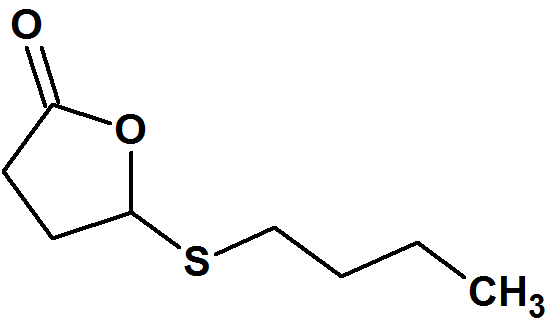 |
| Triethyl phosphate (36) | 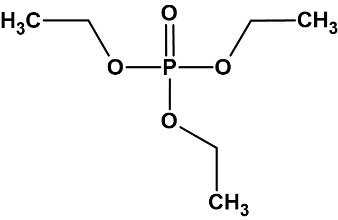 |
| p-nitrophenyl phosphate (37) | 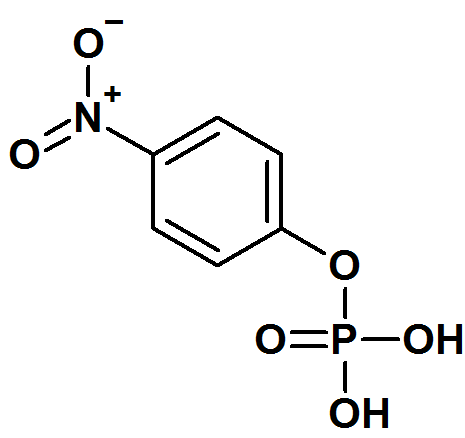 |
| Bis(p-nitrophenyl) phosphate (38) | 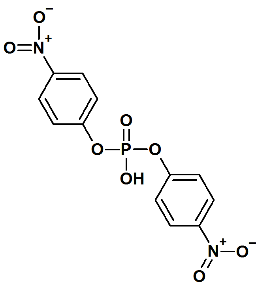 |
| *O*-(4-aminobutyl) *O,O*-diphenyl phosphate (39) | 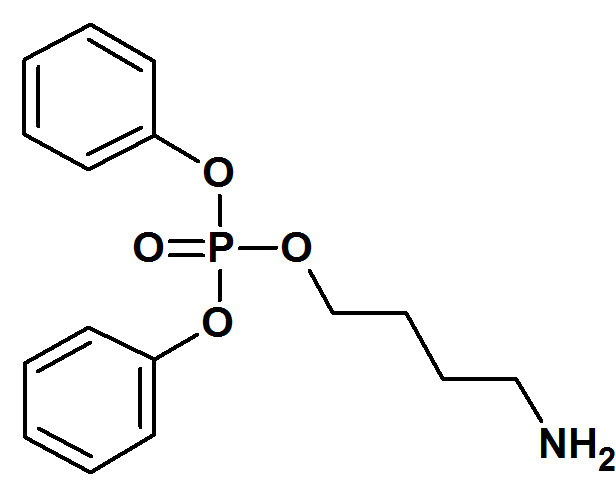 |
| VX  (40) | 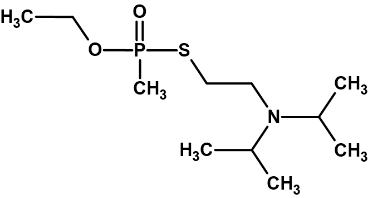 |

**Table S3: The list of isolated strains from the Red Sea and the Mediterranean Sea under various conditions.** RS – Red Sea; MS – Mediterranean Sea, 600 m from the shore; MSM1 - Mediterranean Sea, coastal water mixed with fresh water at location 1; MSM2 - Mediterranean Sea, coastal water mixed with fresh water at location 2. DMAP – dimethyl acetophenone phosphate (compound 10); DMPP – dimethyl phenyl phosphate (compound 9), TMP+TEP – trimethyl phosphate (compound 8) and triethyl phosphate (compound 36); DPAPP – diphenyl acetophenone phosphate (compound 12). Carbon source: Glc – glucose; Suc – succinate; Glyc – glycerol.

| **AOP** | **C source** | **Sampling Location** | **Closest Identified Bacteria**  **(seq ID)** | **Cover**  **(%)** | **Seq. data (16S rRNA identity %)** | **Order** | **Comments** |
| --- | --- | --- | --- | --- | --- | --- | --- |
| DMAP | Glc | RS | - | - | - | - |  |
|  |  | MS | *Cobetia sp.* AM6  (AP021868.1) | 100 | 1395/1395 (100%) | Oceanospirillales | Full size genome sequenced |
|  |  |  | *Celeribacter naphthalenivorans* EMB201  (NR_137260.1) | 100 | 1293/1301 (99.4%) | Rhodobacterales | Partial genome sequenced  Naphthalene-degrading bacterium *(10)* |
|  |  | MSM1 | *Pseudoalteromonas sp.* Xi13  (CP034439.1) | 100 | 1412/1413 (99.9%) | Alteromonadales | Full size genome sequenced |
|  |  | MSM2 | *Celeribacter naphthalenivorans* EMB201  (NR_137260.1) | 100 | 1293/1301 (99.4%) | Rhodobacterales |  |
|  |  |  | *Cobetia sp.* AM6  (AP021868.1) | 100 | 1373/1373 (100%) | Oceanospirillales |  |
|  | Suc | RS | *Vibrio sp.* THAF190c  (CP045338.1) | 100 | 1384/1387 (99.8%) | Vibrionales | Full size genome sequenced isolated from the surface of a PET microplastic particle |
|  |  | MS | *Alteromonas macleodii Sh6* (KP843697.1) | 100 | 1378/1386 (99.4%) | Alteromonadales | Partial genome sequenced |
|  |  | MSM1 | *Phaeobacter sp.* R-52693  (KT185143.1) | 100 | 1313/1313 (100%) | Rhodobacterales | Partial genome sequenced |
|  |  |  | *Marinomonas brasilensis* R-40503  (NR_122094.1) | 100 | 1390/1392 (99.9%) | Oceanospirillales | Partial genome sequenced  Isolated from the coral |
|  |  | MSM2 | *Celeribacter naphthalenivorans* EMB201  (NR_137260.1) | 100 | 1310/1318 (99.4%) | Rhodobacterales |  |
|  |  |  | *Marinobacter sp.* SS1.34  (KC160867.1) | 100 | 1367/1370 (99.8%) | Alteromonadales | Partial genome sequenced  *(11)* |
|  | Glyc | RS | *Vibrio sp.* PaD2.05  (GQ406613.1) | 100 | 1355/1355 (100%) | Vibrionales | Partial genome sequenced  *(12)* |
| DMPP | Glc | RS | *Alteromonas mediterranea* U4  (CP004849.1) | 100 | 1376/1378 (99.9%) | Alteromonadales | Full size genome sequenced *(13)* |
|  |  | MS | - | - | - | - |  |
|  |  | MSM1 | *Labrenzia sp.* MBE 8  (KF724485.1) | 100 | 1350/1350 (100%) | Hyphomicrobiales | Partial genome sequenced |
|  |  | MSM2 | *Ruegeria sp.* InAD-147  (MF401348.1) | 100 | 1311/1313 (99%) | Rhodobacterales | Partial genome sequenced |
|  | Suc | RS | *Alteromonas mediterranea* U4  (CP004849.1) | 100 | 1328/1330 (99.8 %) | Alteromonadales |  |
|  |  |  | *Phaeobacter sp.* R-52693 (KT185143.1) | 100 | 1282/1283 (99.9%) | Rhodobacterales | Partial genome sequenced  *(14)* |
|  |  | MS | *Alteromonas macleodii* Sh6 (KP843697.1) | 99 | 1392/1399 (99.5%) | Alteromonadales |  |
|  |  | MSM1 | *Phaeobacter sp.* R-52693  (KT185143.1) | 100 | 1333/1333 (100%) | Rhodobacterales |  |
|  |  | MSM2 | *Cobetia sp.* AM6  (AP021868.1) | 100 | 1419/1419 (100%) | Oceanospirillales |  |
| TMP+TEP | Glc | RS | *Tateyamaria omphalii* MKT107 (NR_125446.1) | 99 | 1285/1298 (99%) | Rhodobacterales | Partial genome sequenced |
|  |  | MS | *Celeribacter naphthalenivorans* EMB201  (NR_137260.1) | 100 | 1313/1321(99%) | Rhodobacterales | Partial genome sequenced |
|  |  | MSM1 | *Phaeobacter sp.* CfWS2c  (MN099587.1) | 99 | 1300/1315 (98.9%) | Rhodobacterales | Partial genome sequenced  *(15)* |
|  |  |  | *Celeribacter neptunius* strain H 14  (NR_116721.1) | 100 | 1321/1321(100%) | Rhodobacterales | Partial genome sequenced |
|  |  | MSM2 | *Ruegeria sp.* L21-PYE-C37  (KJ188015.1) | 100 | 1341/1341 (100%) | Rhodobacterales |  |
|  | Suc | RS | - | - | - | - |  |
|  |  | MS | *Celeribacter naphthalenivorans* EMB201  (NR_137260.1) | 100 | 1310/1318 (99.4%) | Rhodobacterales |  |
|  |  | MSM1 | *Phaeobacter sp.* CfWS2c  (MN099587.1) | 99 | 1300/1315 (98.9%) | Rhodobacterales |  |
|  |  |  | *Celeribacter neptunius* strain H 14  (NR_116721.1) | 100 | 1313/1313(100%) | Rhodobacterales | Partial genome sequenced  *(16)* |
|  |  | MSM2 | *Celeribacter sp.* KWE30-14  (JQ670713.1) | 100 | 1332/1345  (99%) | Rhodobacterales | Partial genome sequenced |
|  |  |  | *Ruegeria sp.* InAD-147  (MF401348.1) | 100 | 1335/1336 (99.9%) | Rhodobacterales | Partial genome sequenced |
| DPAPP | Glc | RS | *Phaeobacter sp.* R-52693  (KT185143.1) | 100 | 1311/1311 (100%) | Rhodobacterales |  |
|  | Suc | RS | *Phaeobacter sp.* R-52693  (KT185143.1) | 100 | 1285/1285 (100%) | Rhodobacterales |  |

**Table S4: Kinetic characterization of the newly discovered PTE cyclase-PTE from *R. pomeroyi* DSS3 and the metallo-beta lactamase (MBL-PTE) from *Ruegeria sp.* TM1040.**

| **Substrate** | **Cyclase-PTE** | | | **MBL-PTE** | | |
| --- | --- | --- | --- | --- | --- | --- |
|  | **K_M_**  **(mM)** | **k_cat_ (min^-1^)** | **k_cat_/K_M_**  **(M^-1^min^-1^)** | **K_M_ (mM)** | **k_cat_**  **(min^-1^)** | **k_cat_/K_M_**  **(M^-1^min^-1^)** |
| Methyl paraoxon (24) | 1.10±0.11 | 12271±602 | 1.02x10^7^ | 1.21±0.11 | 71.20±2.61 | 5.88x10^4^ |
| Paraoxon (25) | 0.68±0.05 | 486±15 | 7.15x10^5^ | 0.90±0.08 | 4.88±0.15 | 5.42x10^3^ |
| Phenyl paraoxon (26) | NA | NA | NA | 0.10±0.01 | 109±7 | 1.09x10^6^ |
| Parathion (30) | NA | NA | NA | NA | NA | NA |
| Methyl parathion (31) | 0.26±0.11 | 0.88±0.10 | 3.40x10^3^ |  |  | 197.2 |
| GD (32) | ND | ND | Fast: 6.74±1.00x10^7^  Slow: 0.80±0.13x10^7^ | ND | ND | Fast:  2.75±0.77x10^5^  Slow: 0.44±0.15x10^5^ |
| GF (33) | ND | ND | 9.55±0.50x10^7^ | ND | ND | 8.15±0.49x10^4^ |
| Isatin (34) | 0.18±0.04 | 261±16 | 1.46x10^6^ | NA | NA | NA |
| TBBL (35) | 0.09±0.03 | 874±170 | 9.53x10^6^ | 0.84±0.24 | 29.85±2.75 | 3.55x10^4^ |
| Dimethyl acetophenone phosphate (10) | 1.53±0.14 | 209±10 | 1.36x10^5^ | 0.81±0.17 | 10.01±0.99 | 1.24x10^4^ |

NA, not active under the experimental conditions employed

ND, not determined

**Table S5. Co-expression of chaperones with cyclase.** Cells were grown in LB medium supplemented with 0.5 mg/ml arabinose, 0.1 % glucose and 1mM ZnCl_2_ at 37°C. At OD=0.6 1mM IPTG was added and expression continued at 30°C or 18°C overnight. Activity was measured in the cell lysate with 1 mM methyl paraoxon (compound 24).

| **Plasmids** | **Promoter (chaperone genes)** | **Expression Temperature (°C)** | **Activity (µM/min)** |
| --- | --- | --- | --- |
| pACYC +pET21-cyclase  (empty vector - control) | *araBp* | 30 | 0.4 |
|  |  | 18 | 1.5 |
| pGro7 + pET21-cyclase | *araBp* | 30 | 24.4 |
|  | *(groES-groEL)* | 18 | 310.1 |
| pKJE7 + pET21-cyclase | *araBp* | 30 | 0.02 |
|  | *(groES-groEL- dnaK-dnaJ-grpE)* | 18 | 0.1 |
| pG-KJE8 + pET21-cyclase | *araBp* | 30 | 0.6 |
|  | *(dnaK-dnaJ-grpE)* | 18 | 28.7 |
| pTF16 + pET21-cyclase | *araBp* | 30 | 0.2 |
|  | *(tig)* | 18 | 0.2 |
